# EXO1 Facilitates MiDAS and Prevents Genome Instability and Cell Death in Ewing Sarcoma

**DOI:** 10.64898/2026.06.05.730187

**Authors:** Joaquín Olmedo-Pelayo, Laura Lobo-Selma, Daniel Delgado-Bellido, Carmen Jordan-Pérez, Paula Gilabert-Prieto, Marco Pérez, Florian H. Geyer, Martha J. Carreño-Gonzalez, Javier Alonso, Li Zheng, Binghui Shen, Thomas G. P. Grünewald, Fernando Gómez-Herreros, E. de Álava

## Abstract

Ewing sarcoma (EwS) is an aggressive malignancy driven by EWSR1::ETS fusions, predominantly EWSR1::FLI1. Previous efforts using both direct and indirect approaches to target these chimeric oncoproteins have yielded limited clinical benefit. Although EWSR1::FLI1 is a well-known source of replication stress and genome instability, targeting DNA damage response (DDR) factors that mitigate these effects remain poorly understood. Here, we identified a marked dependency of EwS cells on exonuclease 1 (EXO1). We demonstrate that EXO1 is essential for EwS cell survival and tumor growth, highlighting its potential as a novel therapeutic target. Intriguingly, we unveil that EXO1 loss impairs mitotic DNA synthesis (MiDAS), promoting EWSR1::FLI1-associated genome instability and cell death. Collectively, our results support the idea that targeting DDR factors, which counteract replication stress and/or DNA damage induced by fusion oncoproteins, represents a promising therapeutic option for EwS.

## INTRODUCTION

Fusion genes resulting from chromosomal rearrangements are common driver mutations in hematological malignancies and childhood sarcomas^1^, representing highly relevant targets for precision medicine. Based on their molecular function, chimeric oncoproteins encoded by fusion genes can be classified into deregulated kinases, such as BCR::ABL in chronic myeloid leukemia^2^; or aberrant transcription factors, such as EWSR1::ETS fusions in Ewing sarcoma (EwS)^3^. Although several drugs targeting fusion kinases have been successfully translated into clinical practice^4^, chimeric transcription factors have long represented a therapeutic challenge and were historically considered “undruggable”^5^.

EwS is a highly aggressive cancer that primarily affects children, adolescents, and young adults^6^. Current multimodal treatment, including intense chemotherapy, radiotherapy, and surgical resection, has substantially improved survival in patients with localized disease^7^. However, prognosis for patients with metastatic or relapsed disease remains dismal^7,8^. Moreover, acute and long-term treatment-associated toxicity can significantly affect the quality of life of survivors^9^. Therefore, the development of more effective and less toxic targeted therapeutic options for EwS is urgently needed.

Genetically, EwS is characterized by pathognomonic chromosomal translocations that generate gene fusions involving *EWSR1* and an ETS gene family member, mainly *FLI1* (∼85% of the cases)^3^. The EWSR1::FLI1 oncoprotein functions as an abnormal transcription factor that plays a key role in the malignant transformation of EwS by reprogramming the cellular transcriptome^10,11^. Although EWSR1::FLI1 could be considered an outstanding target for EwS therapy, direct targeting remains challenging due to several reasons, including its nuclear localization, its activity as a transcription factor, and the ubiquitous expression of *EWSR1* and *FLI1* genes in non-tumoral tissues^9^. By contrast, indirect targeting of EWSR1::FLI1, mainly focused on inhibiting its transcriptional cofactors and downstream targets, has been extensively explored^12,13^. Unfortunately, although most of these strategies have shown promising results in preclinical studies, their impact on EwS therapy has been limited.

Prior efforts to generate cellular and animal models of EwS have demonstrated that ectopic expression of the *EWSR1::FLI1* oncogene is highly toxic to most cell types^14–18^. Furthermore, elevated levels of EWSR1::FLI1 in EwS cells lead to DNA damage and cell death^19^, which may be explained by the ability of the fusion protein to induce replication stress^20–22^, a well-known source of genome instability^23^. Nevertheless, the possible role of DNA damage response (DDR) factors in mitigating EWSR1::FLI1-driven replication stress and/or preventing genome instability remains almost unexplored.

In this study, we identified exonuclease 1 (EXO1) as a clinically relevant targetable candidate. We demonstrated that EWSR1::FLI1 renders EwS cells vulnerable to EXO1 depletion and inhibition. In fact, EXO1 loss promotes EWSR1::FLI1-induced genome instability in mitosis by disrupting mitotic DNA synthesis (MiDAS). Overall, our findings suggest that targeting DDR factors that support tolerance of EwS cells to EWSR1::FLI1 represents a promising approach for the development of novel therapeutic strategies.

## RESULTS

### *EXO1* is overexpressed in EwS and correlates with unfavourable patient outcome

To identify potential therapeutic targets in EwS associated with genome repair and maintenance, we interrogated available microarray expression data from 166 clinically annotated EwS tumors^24^. We assessed the significance of the association between 201 DDR factors and overall survival, stratifying samples into tertiles based on gene expression levels. Among DDR factors significantly associated with EwS outcome (Bonferroni-adjusted P < 0.05), *RAD54L*, which is overexpressed in several cancer models^25^, achieved the highest score (Fig. 1A). However, we focused on EXO1, a 5’-3’ double-stranded DNA exonuclease that plays a crucial role in multiple DNA metabolic processes such as DNA replication and repair^26^. Importantly, among top hits, *EXO1* gene was the only one located on the long arm of chromosome 1 (chr1q), whose gain has been previously associated to an aggressive and highly proliferative form of EwS^27,28^. Importantly, the correlation between elevated *EXO1* expression and reduced overall survival observed in this cohort (Fig. 1B), was validated at protein levels in an independent cohort of 76 EwS samples (Fig. 1C, Supplementary Fig. 1A). Comprehensive exploration of public datasets showed that *EXO1* is highly expressed in EwS compared to normal tissues and other tumoral cell lines and tumor samples (Supplementary Fig. 1B & 1C, Fig. 1D). This observation was corroborated by immunohistochemistry in a tissue microarray comprising several sarcoma subtypes (Fig. 1E). Given that chr1q is gained in around 18% of EwS patients^29^, we investigated whether it could contribute to the inter-tumoral heterogeneity of *EXO1* expression among EwS tumours. Interestingly, *EXO1* expression significantly correlated with chr1q copy number in EwS cell lines (Supplementary Fig. 1D). Due to the absence of public datasets integrating both copy number and expression data, we inferred chr1q status of the previous 166 patient samples using an EwS-specific chr1q gain expression signature (see methods for details). Notably, chr1qG “high” patients exhibited reduced overall survival, in agreement with previous reports (Supplementary Fig. 1E)^28,29^. More importantly, *EXO1* expression levels were significantly higher in chr1qG “high” patient samples (Fig. 1F). Altogether, our findings indicate that *EXO1* is highly but heterogeneously expressed in EwS and is significantly associated with poor patient outcomes.

**Figure 1.**
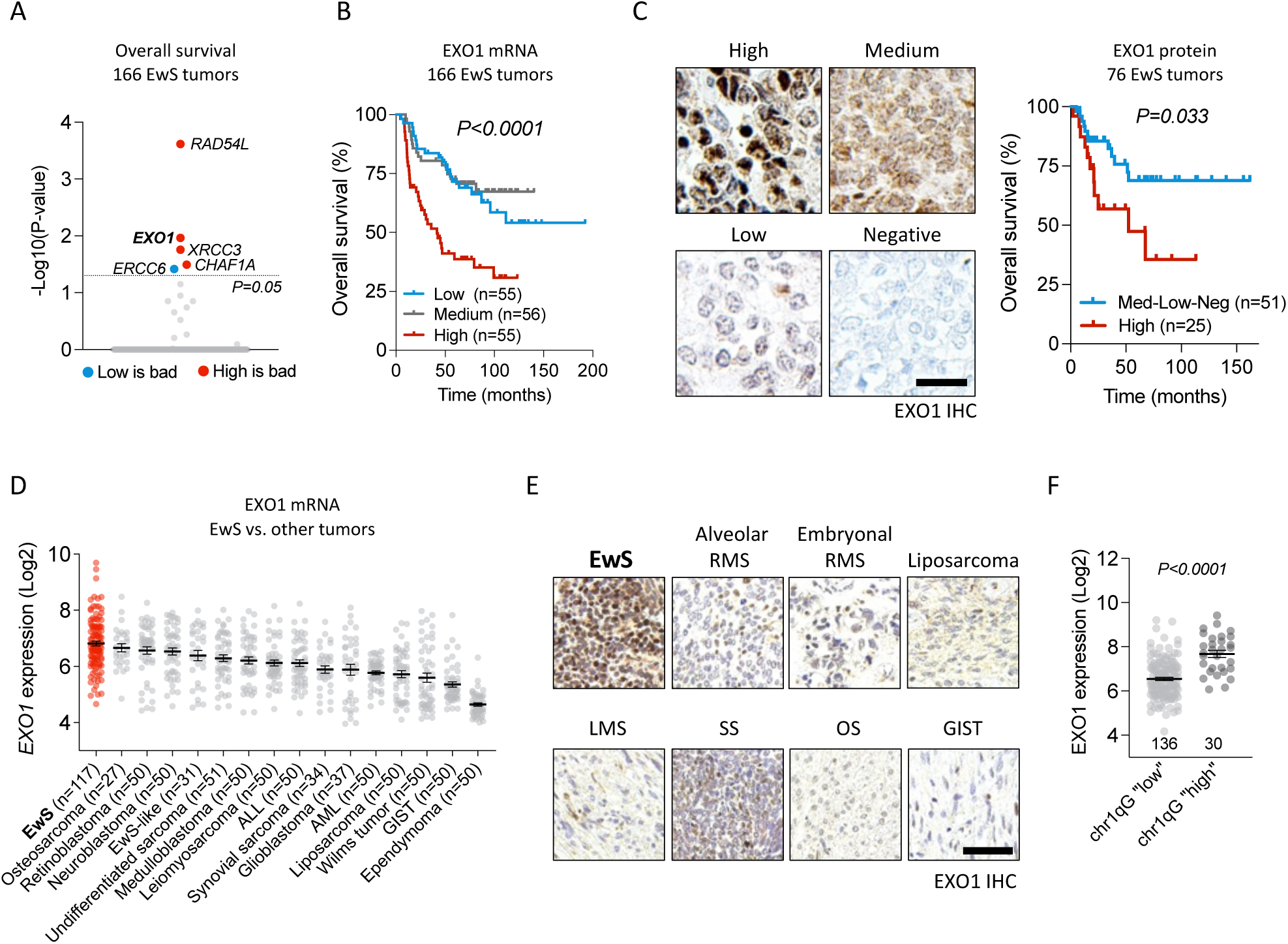
*EXO1* is overexpressed in EwS and correlates with unfavourable patient outcome. **(A)** Analysis of the association between 201 DDR genes and EwS overall survival among 166 clinically annotated EwS transcriptomes. *P* value was determined by the Mantel-Cox test and corrected by the Bonferroni method. **(B)** Kaplan-Meier survival analysis of 166 clinically annotated EwS transcriptomes, stratified into tertiles by EXO1 expression level. *P* value was determined by Mantel-Cox test. **(C)** Kaplan-Meier survival analysis of 76 EwS samples stratified according to EXO1 staining score (intensity x extension). *P* value was determined by Mantel-Cox test. *Left*, representative images of EXO1 IHC. Scale bar: 50 µm. **(D)** *EXO1* expression levels (mean ± SEM) in EwS, other sarcomas, and pediatric tumors. **(E)** Evaluation of EXO1 protein levels in a tissue microarray comprising several sarcoma subtypes by IHC (RMS, rhabdomyosarcoma; LMS, leiomyosarcoma; SS, synovial sarcoma; OS, osteosarcoma; GIST, gastrointestinal stromal tumor). Scale bar: 50 µm. **(F)** *EXO1* expression levels (mean ± SEM) between EwS transcriptomes with “high” and “low” chr1q expression signature. *P* value was determined by two-tailed unpaired t-test.

### EWSR1::FLI1 confers a dependency on EXO1 in EwS cells

To assess the potential dependency of EwS cell survival on EXO1, we carried out clonogenic assays in two EwS cell lines with low (A-673) and moderate–high (RD-ES) *EXO1* expression levels (Supplementary Fig. 2A). Additionally, the osteosarcoma cell line U2-OS, with similar *EXO1* expression to RD-ES, was assessed (Supplementary Fig. 2B). Interestingly, siRNA-based *EXO1* silencing strongly reduced the clonogenic capacity of EwS cell lines in comparison to U2-OS (Fig. 2A, Supplementary Fig. 2C). To confirm the dependency of EwS cells on EXO1 in an isogenic background, we generated a RD-ES cellular model carrying a doxycycline (DOX)-inducible shRNA against *EXO1* (RD-ES/TR/shEXO1), and a non-targeting control (RD-ES/TR/shCtrl). Incubation of these cells with DOX efficiently reduced EXO1 at mRNA and protein levels (Fig. 2B, Supplementary Fig. 2D). In agreement with our previous findings, DOX-mediated EXO1 depletion significantly impaired cellular clonogenicity, and anchorage-independent cellular growth (Fig. 2C & 2D). Notably, growth inhibition was associated with the induction of apoptosis, as evidenced by an increased percentage of AnnexinV-positive cells and elevated levels of the apoptotic marker cleaved caspase-3 upon *EXO1* downregulation (Fig. 2E & 2F). To directly evaluate the effect of EXO1 loss on tumor growth, we generated a xenograft model by injecting RD-ES/TR/shEXO1 cells subcutaneously in immunocompromised mice. Importantly, DOX-mediated EXO1 depletion significantly impaired tumor growth (Fig. 2G). Collectively, these results highlight EXO1 as an essential factor for EwS survival. Next, we investigated the contribution of EWSR1::FLI1 fusion protein to the EwS dependency on EXO1. We employed a previously established HeLa model (HeLa EF), which expresses the *EWSR1::FLI1* oncogene under the control of a DOX-inducible promoter^30^. As previously shown, ectopic *EWSR1::FLI1* expression resulted in a significant reduction of cellular survival (Fig. 2H). More importantly, although *EXO1* silencing reduced clonogenicity of control (DOX-) cells, this effect was much stronger in cells overexpressing the fusion oncogene (Fig. 2H). Treatment with C73, the first-in-class EXO1 inhibitor^31^ confirmed the dependency of EwS cells on EXO1 (Fig. 2I). Altogether, these results uncover a marked vulnerability of EwS cells to EXO1 perturbations, which is driven by the EWSR1::FLI1 oncoprotein.

**Figure 2.**
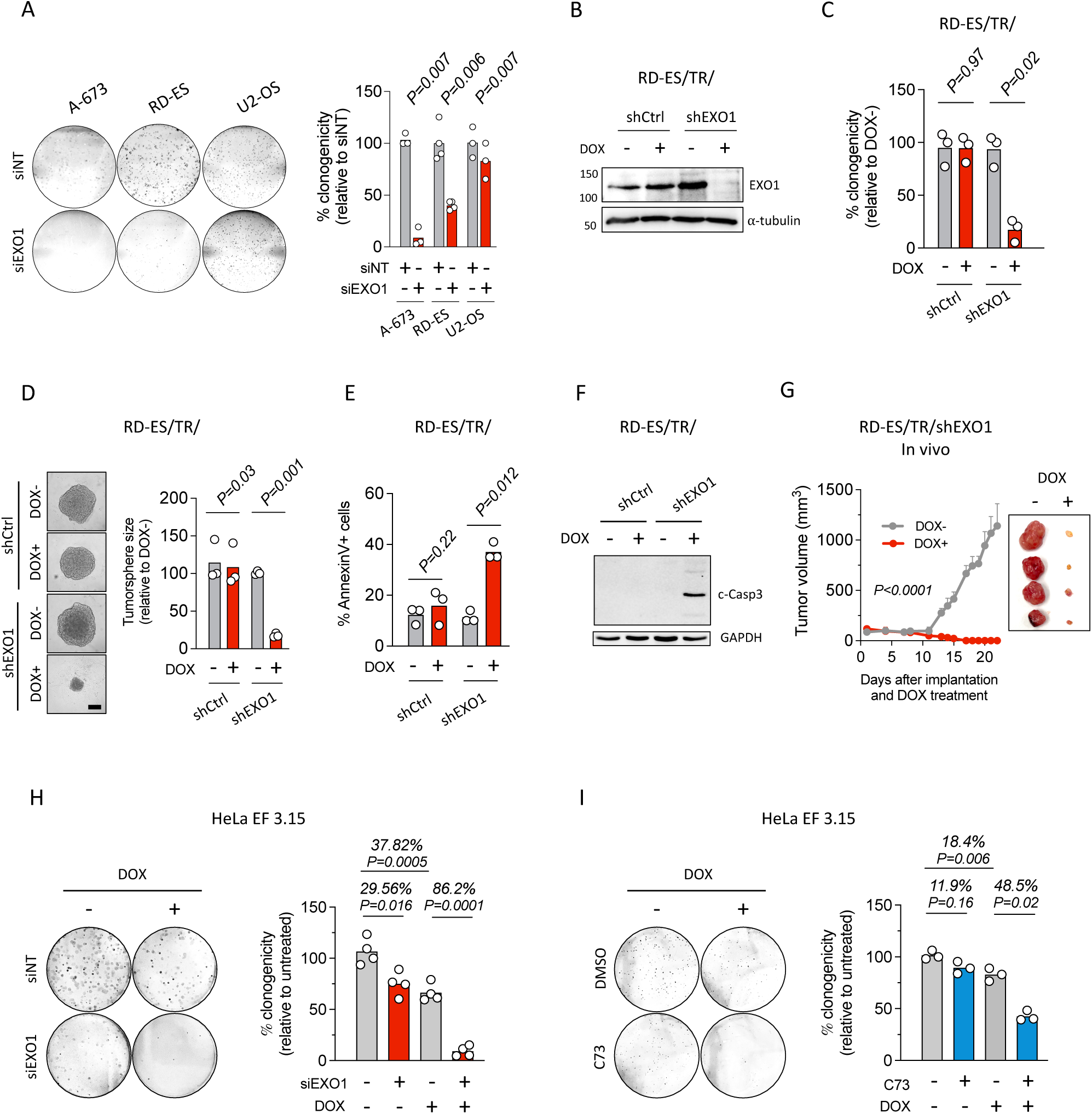
EWSR1::FLI1 renders EwS dependency on EXO1. **(A)** Analysis of the effect of siRNA-based *EXO1* downregulation on the clonogenic capacity of A-673, RD-ES and U2-OS cells. *Left,* representative images. *Right,* data represent the mean number of colonies (relative to siNT); n≥3 independent experiments. **(B)** Evaluation of EXO1 protein levels in RD-ES/TR/shEXO1 (and shCtrl) cells after 72 h of DOX incubation by WB. Molecular weight in kDa. ⍺-tubulin: loading control. **(C)** Analysis of the effect of *EXO1* downregulation on cellular clonogenicity. Cells were grown with or without DOX for 10-12 days. Data represent the mean number of colonies (relative to DOX-); n=3 independent experiments. **(D)** Evaluation of the effect of *EXO1* silencing on anchorage-independent cellular growth. Cells were grown in ultra-low attachment plates with or without DOX for 6 days*. Left*, representative images. Scale bar: 500 µm. *Right,* data represent the mean of the tumor sphere size (relative to DOX-); n=3 independent experiments. **(E)** Study of the induction of apoptosis after *EXO1* silencing by Annexin V FACS. Cells were incubated with DOX for 4 days. Data represent the mean of the percentage of Annexin V-positive cells; n=3 independent experiments. **(F)** Similar to (E) by WB for the evaluation of cleaved caspase-3. Molecular weight in kDa. GAPDH: loading control**. (G)** Analysis of the effect of EXO1 depletion on tumor growth. RD-ES/TR/shEXO1 cells were subcutaneously injected in immunocompromised mice, and DOX was administered at 2 mg/ml via drinking water for 20 days. *Left,* data represent the mean (+SEM) of tumor volume along DOX treatment; n=4 tumors per condition. *Right,* image of tumors at the end of treatment. **(H)** Effect of *EXO1* silencing in the clonogenic capacity of HeLa EF cells. When indicated, cells were incubated with DOX for 72 h and transfected with indicated siRNAs for 48 h. After that, cells were seeded at low density and grown for 7 days. *Left,* representative images. *Right,* data represent the mean number of colonies (relative to DOX-siNT); n=4 independent experiments. **(I)** Similar to (H) after incubation with DOX for 72 h and/or 2.5 µM of C73 for 48 h; n=3 independent experiments. *P* value was determined by two-tailed paired t-test in panels A, C, D, E, H and I; and by two-way ANOVA in panel G.

### EXO1 prevents DNA damage and chromosomal instability in EwS

Given the role of EXO1 in several DNA repair pathways, we investigated whether the deleterious effect of EXO1 loss on EwS viability was associated with the induction of double-strand breaks (DSBs), the most cytotoxic form of DNA damage. To explore this possibility, we evaluated phospho-H2AX (Ser139; hereafter γH2AX), a well-established surrogate marker of DSBs^32^. Interestingly, *EXO1* silencing increased γH2AX levels in EwS cells, with no detectable effect in the U2-OS cell line (Fig. 3A & 3B). These findings were further validated by EXO1 inhibition (Supplementary Fig 3A). EXO1 plays a key role in DNA end resection, a critical step in the repair of DSBs through homologous recombination^33^. In addition, it has recently been shown that EXO1 is an essential factor in *BRCA1*-mutated cells for single-strand annealing (SSA) repair. Within this cellular context, EXO1 loss induces DNA lesions in S-phase^34^. Considering that EWSR1::FLI1 impairs BRCA1 activity in EwS^22^, we speculated that EXO1 depletion may cause genome instability by disrupting SSA repair. To test this possibility, we measured γH2AX in S-phase population, which was determined by exposing cells to a short pulse of 5-Ethylnyl-2’-deoxyuridine (EdU), an analogue of thymidine incorporated during DNA synthesis. Strikingly, DOX-mediated *EXO1* silencing significantly increased γH2AX levels in EdU-negative cells, whereas no differences were observed in the EdU-positive (S-phase) population (Fig. 3C). To corroborate this observation, we evaluated γH2AX distribution along the cell cycle by flow cytometry. Etoposide, a DNA topoisomerase II poison, was used as a positive control. In agreement with our previous finding, EXO1 depletion significantly accumulated DSBs in G1-phase cells, while no differences were detected within S or G2 phases (Fig. 3D). In fact, *EXO1* silencing was associated with a subtle increase of G1-phase population (Supplementary Fig. 3B). Altogether, our results indicate that EXO1 prevents DNA damage accumulation in G1-phase EwS cells.

**Figure 3.**
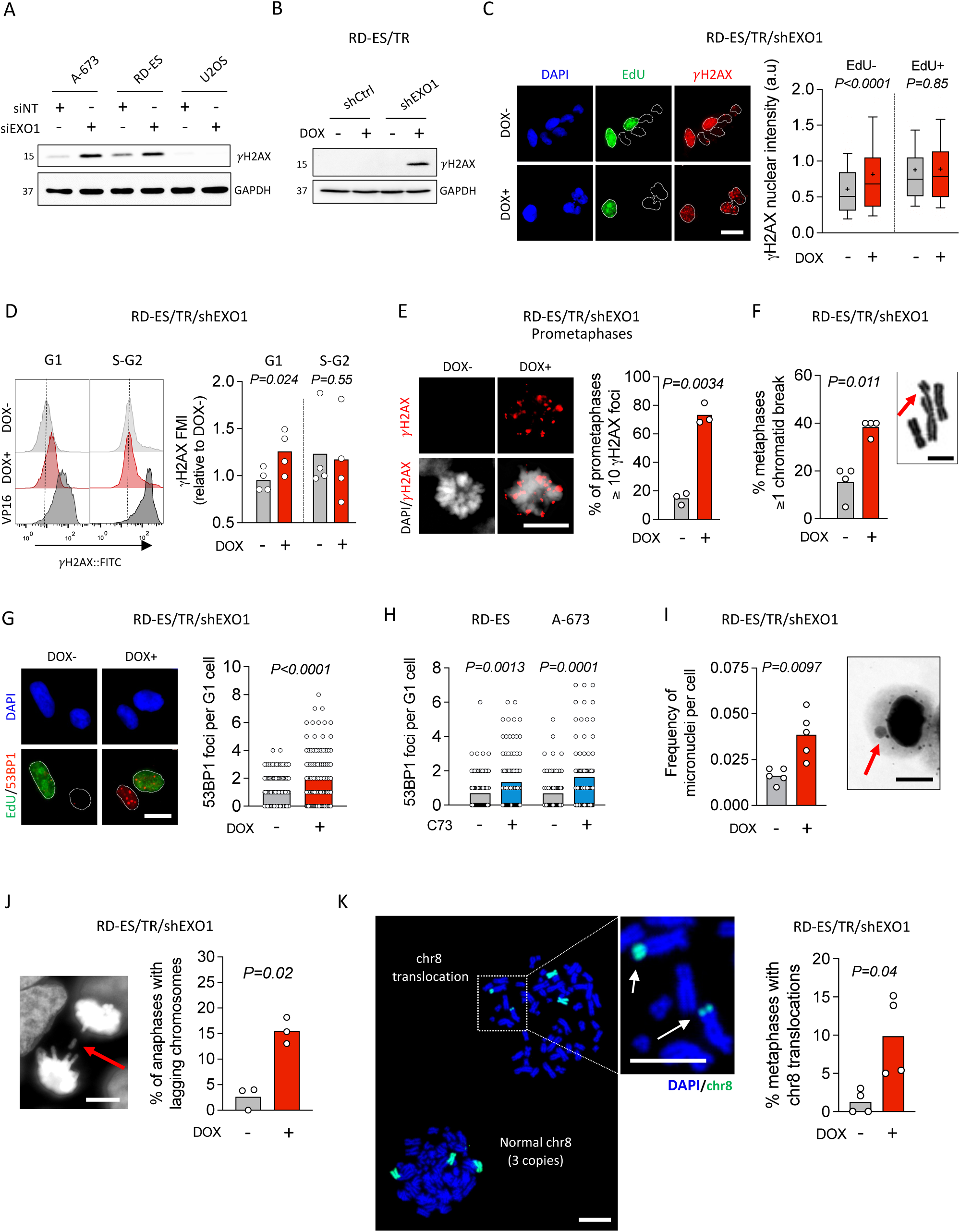
EXO1 prevents DNA damage and chromosomal instability in EwS. **(A)** Analysis of the effect of *EXO1* silencing on the induction of DSBs by !H2AX WB. Cells were transfected with the indicated siRNAs for 72 h. Molecular weight in kDa. GAPDH: loading control. **(B)** Similar to (A) in RD-ES/TR/shEXO1 (and shCtrl) cells after 48 h of DOX incubation. **(C)** Analysis of DSBs accumulation after EXO1 depletion by !H2AX IF. When indicated, RD-ES/TR/shEXO1 cells were incubated with DOX for 48 h. EdU (10 µM) was added to the culture media for the last 20 min of DOX incubation. *Left,* representative images. DAPI counterstain. Scale bar: 20 µm. *Right,* data represent the mean (+)(±10-90 percentile) of nuclear !H2AX intensity in EdU– and EdU+ cells; n=3 independent experiments (∼100 cells were analyzed per replicate). **(D)** Evaluation of !H2AX distribution along the cell cycle after *EXO1* knockdown. When indicated, cells were incubated with DOX for 48 h or with 20 µM of etoposide for 1 h. *Left,* histogram of !H2AX signal in G1 and S-G2 phase populations (determined by PI). *Right,* data represent the mean of !H2AX intensity (relative to DOX-); n=4 independent experiments. **(E)** Evaluation of the induction of DSBs in prometaphases upon EXO1 depletion by !H2AX IF. When indicated, cells were incubated with DOX for 48 h. *Left,* representative images. DAPI counterstain. Scale bar: 10 µm. *Right,* data represent the mean percentage of prometaphases with 10 or more !H2AX foci; n=3 independent experiments. **(F)** Analysis of the effect of *EXO1* downregulation on the induction of chromatid breaks. Cells were incubated with DOX for 48 h. *Left,* data represent the mean of metaphases with 1 or more chromatid breaks; n=4 independent experiments. *Right,* representative image. Red arrow indicates a chromatid break. Giemsa staining. Scale bar: 5 µm. **(G)** Evaluation of the effect of EXO1 depletion on the formation of 53BP1-NBs. When indicated, cells were incubated with DOX for 48 h. EdU (10 µM) was added to the culture media for the last 15 min of DOX incubation. *Left,* representative images. DAPI counterstain. Scale bar: 20 µm. *Right,* data represent the mean of 53BP1-NBs per G1-phase cell; n=3 independent experiments (∼30 cells were analyzed per replicate). **(H)** Similar to (G) after treatment with 2.5 µM of C73 for 72 h; n=3 independent experiments. **(I)** Effect of *EXO1* silencing on micronuclei formation. Cells were incubated with DOX for 48 h. *Left,* data represent the mean of the frequency of micronuclei per cell; n=5 independent experiments. *Right*, representative image. Red arrow indicates a micronucleus. Giemsa staining. Scale bar: 50 µm. **(J)** Effect of EXO1 depletion on the levels of lagging chromosomes. Cells were incubated with DOX for 48 h. *Left,* representative image. Red arrow indicates a lagging chromosome. DAPI counterstain. Scale bar: 50 µm. *Right,* data represent the mean of anaphases showing lagging chromosomes; n=3 independent experiments. **(K)** Analysis of the effect of EXO1 depletion on the formation of chromosomal translocations by chr8 FISH. Cells were incubated with DOX for 72 h. *Left*, representative image. White arrows indicate chr8 rearrangements. DAPI counterstain. Scale bar: 50 µm. *Right,* data represent the percentage of metaphases showing chr8 translocations. *P* value was determined by two-tailed unpaired t-test in panels C, G and H; and paired t-test in panels D, E, F, I, J and K.

Given that EXO1 is degraded in G1 phase to prevent DNA end resection in the absence of a homologous DNA template^35^, we hypothesized that EXO1 loss could induce DSBs during previous mitosis. To evaluate this possibility, we analyzed γH2AX in prometaphases. Interestingly, EXO1 depletion significantly increased the percentage of cells exhibiting high levels of DNA damage (10 or more γH2AX foci) (Fig. 3E). In accordance, flow cytometry using the mitotic marker phospho-histone 3 (Ser10; hereafter pH3S10) showed that EXO1 silencing and inhibition increased γH2AX signal in mitotic (pH3S10+) cells (Supplementary Fig. 3C & 3D). Given that γH2AX is a surrogate marker of DSBs, we directly measured chromosomal breaks in metaphase spreads. Accurately, we observed a higher frequency of metaphases with chromatid breaks, an indicator of post-replicative DSBs, in cells lacking EXO1 (Fig. 3F). DSB repair pathways are not efficiently activated in mitosis, and cells progress to the next G1 phase harbouring 53BP1-marked lesions, forming the so-called 53BP1 nuclear bodies (53BP1-NBs)^36^. Strikingly, EXO1 depletion and inhibition increased the levels of 53BP1-NBs in G1-phase cells, which were differentiated by their size and the lack of EdU incorporation (Fig. 3G & 3H). These data are consistent with a substantial fraction of EXO1-loss-associated damage becoming evident during mitosis and being transmitted to the next G1 phase.

Given the well-established relationship between mitotic DNA damage and chromosomal instability, we further investigated whether EXO1 depletion promotes chromosomal aberrations in EwS cells. Interestingly, *EXO1* silencing increased the frequency of micronuclei, a well-established hallmark of chromosomal instability^37^ (Fig. 3I). In agreement, EXO1 depletion and inhibition resulted in elevated levels of lagging chromosomes during anaphase, confirming that EXO1 loss triggers mitotic errors in EwS cells (Fig. 3J, Supplementary Fig. 3E). Moreover, FISH for whole chr8 showed that EXO1 silencing and inhibition increased the percentage of metaphases with a gain of Chr8 (4 copies)(Supplementary Fig. 3F & 3G), and the frequency of translocations involving this chromosome (Fig. 3K, Supplementary Fig. 3H). Collectively, these findings demonstrate that EXO1 prevents chromosomal instability in EwS cells.

### EXO1 loss promotes EWSR1::FLI1-associated mitotic genome instability by disrupting MiDAS repair

Next, we investigated the role of EWSR1::FLI1 in the genome instability associated to EXO1 depletion. Notably, EXO1 loss promoted higher levels of γH2AX in HeLa EF cells expressing ectopically the *EWSR1::FLI1* oncogene compared to the control (DOX-) counterparts (Fig. 4A). Consistently, similar results were obtained upon treatment with C73 inhibitor (Supplementary Fig. 4A). To corroborate these findings, we employed a pre-established A-673 cellular model carrying a DOX-inducible shRNA against *EWSR1::FLI1* (A-673/TR/shEF)(Supplementary Fig. 4B)^38^. Interestingly, DOX-mediated *EWSR1::FLI1* downregulation significantly reduced the levels of γH2AX induced by *EXO1* silencing (Fig. 4B), confirming that EXO1 loss promotes EWSR1::FLI1-associated DNA damage. Additionally, we analysed whether EXO1 was also crucial for preventing genome instability in cells harbouring distinct fusion oncogenes. Importantly, in comparison to RD-ES, EXO1 inhibition did not increase γH2AX levels in RM-82 (EwS; *EWSR1::ERG*), JN-DSRCT (desmoplastic small round cell tumor; *EWSR1::WT1*) and RH-30 (alveolar rhabdomyosarcoma; *PAX3::FOXO1*) cell lines, suggesting a specific functional interaction between EXO1 and EWSR1::FLI1 (Supplementary Fig. 4C). More importantly, *EXO1* downregulation significantly increased the frequency of 53BP1-NBs in HeLa EF cells overexpressing the fusion oncogene (Fig. 4C), indicating that EXO1 loss leads to EWSR1::FLI1-induced genome instability in mitosis.

**Figure 4.**
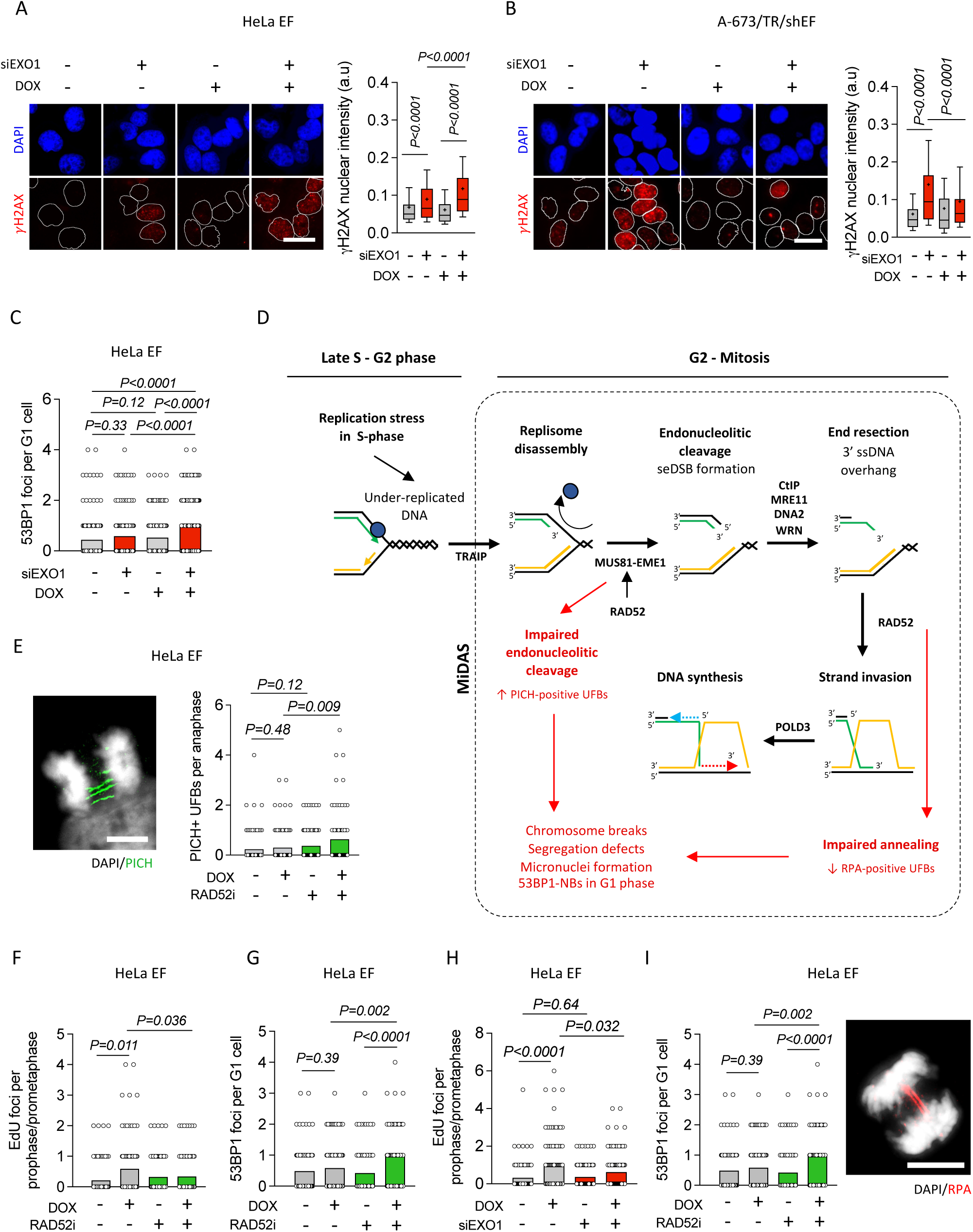
EXO1 loss promotes EWSR1::FLI1-associated mitotic genome instability by disrupting MiDAS repair. **(A)** Analysis of the relationship between EXO1 and EWSR1::FLI1 in the induction of DNA damage by !H2AX IF. When indicated, RD-ES/TR/shEXO1 cells were incubated with DOX for 72 h and transfected with siNT or siEXO1 for 48 h. *Left,* representative images. DAPI counterstain. Scale bar: 20 µm. *Right,* data represent the mean (+)(±10-90 percentile) of nuclear !H2AX intensity; n=3 independent experiments (150 cells were analyzed per replicate). **(B)** Similar to (A) in A-673/TR/shEF cells after incubation with DOX and transfection with siNT or siEXO1 for 48 h; n=3 independent experiments (140 cells were analyzed per replicate). **(C)** Evaluation of the relationship between EXO1 and EWSR1::FLI1 in the induction of 53BP1-NBs. HeLa EF cells were incubated with DOX for 72 h and transfected with siNT or siEXO1 for 48 h. Data represent the mean number of 53BP1-NBs per G1-phase cell; n=3 independent experiments (∼40 cells were analyzed per replicate). **(D)** Scheme of MiDAS repair pathway. **(E)** Evaluation of the effect of EWSR1::FLI1 on the levels of PICH-coated UFBs. When indicated, HeLa EF cells were incubated with DOX for 72 h and/or 20 µM of RAD52 inhibitor AICAR for 1 h. *Left,* representative image. DAPI counterstain. Scale bar: 20 µm. *Right*, data represent the mean number of PICH-positive UFBs per anaphase; n=3 independent experiments (∼30 cells were analyzed per replicate). **(F)** Evaluation of MiDAS activity in HeLa EF cells upon *EWSR1::FLI1* ectopic expression. Cells were treated as indicated in (E). EdU (10 µM) was added to the culture media for the last 20 min of DOX incubation. Data represent the mean number of EdU foci per prophase/prometaphase; n=3 independent experiments (∼30 cells were analyzed per replicate). **(G)** Evaluation of the induction of 53BP1-NBs in HeLa EF cells upon *EWSR1::FLI1* ectopic expression. Cells were treated as in (E). Data represent the mean of 53BP1-NBs per G1-phase cell; n=3 independent experiments (∼30 cells were analyzed per replicate). **(H)** Evaluation of the effect of *EXO1* silencing on EWSR1::FLI1-induced MiDAS repair. When indicated, HeLa EF cells were incubated with DOX for 72 h and transfected with siNT or siEXO1 for 48 h. Data represent the mean number of EdU foci per prophase/prometaphase; n=3 independent experiments. **(I)** Study of the effect of *EXO1* downregulation on EWSR1::FLI1-induced RPA-coated UFBs. *Left,* data represent the mean number of RPA-positive UFBs per anaphase; n=3 independent experiments (∼30 cells were analyzed per replicate). *Right*, representative image. DAPI counterstain. Scale bar: 20 µm. *P* value was determined by two-tailed unpaired t-test.

Replication stress is recognized as a major driver of mitotic genome instability^39^. In fact, reduced replication fork speed results in under-replicated DNA in mitosis, which is processed by MiDAS repair pathway (Fig. 4D)^40^. Given that EWSR1::FLI1 is a well-known source of replication stress, we hypothesized that EXO1 may be required during MiDAS to process late replication intermediates induced by the fusion protein. To test this possibility, we first evaluated whether EWSR1::FLI1 led to the accumulation of under-replicated DNA in mitosis by assessing PICH-coated ultrafine bridges (UFBs) in anaphase^41^. Notably, *EWSR1::FLI1* overexpression *per se*, did not alter the levels of PICH-positive UFBs (Fig. 4E). However, PICH-positive UFBs were accumulated in *EWSR1::FLI1*-overexpressing cells upon inhibition of RAD52 (Fig. 4E), consistent with its role in promoting MUS81-mediated cleavage of late replication intermediates^42^. In contrast, no significant differences were observed upon RAD51 inhibition (Supplementary Fig. 4D), suggesting that EWSR1::FLI1-induced late replication intermediates are processed in a RAD52-dependent manner. Thus, we analysed whether EWSR1::FLI1 activates MiDAS repair by assessing EdU incorporation in mitosis, as a marker of DNA synthesis (Supplementary Fig. 4E). Importantly, *EWSR1::FLI1* overexpression resulted in a significant increase of EdU foci (Fig. 4F). More importantly, RAD52 inhibition strongly reduced EWSR1::FLI1-associated EdU incorporation (Fig. 4F), indicating that EWSR1::FLI1 activates MiDAS repair. Furthermore, RAD52 inhibition led to an additional accumulation of 53BP1-NBs in *EWSR1::FLI1* overexpressing cells (Fig. 4G), suggesting that MiDAS repair prevents EWSR1::FLI1-associated genome instability in mitosis.

Finally, we examined whether EXO1 participates in MiDAS repair facilitating the resolution of late replication intermediates induced by EWSR1::FLI1. Interestingly, EXO1 silencing caused a significant reduction in the frequency of EdU foci induced by *EWSR1::FLI1* expression (Fig. 4H), indicating that EXO1 loss impairs MiDAS repair. However, in contrast to the inhibition of RAD52, EXO1 depletion did not affect the levels of PICH-positive UFBs (Supplementary Fig. 4F), suggesting that EXO1 could act downstream MUS81-dependent endonucleolitic cleavage. To evaluate this possibility, we assessed RPA-positive UFBs, which define recombination intermediates^43^. Importantly, *EXO1* silencing significantly reduced the frequency of EWSR1::FLI1-induced RPA-coated UFBs (Fig. 4I), indicating a potential contribution of EXO1 in facilitating strand invasion. Together, these results support a role for EXO1 in the efficient MiDAS-associated processing of EWSR1::FLI1-induced late replication intermediates, acting downstream of RAD52/MUS81-dependent initiation.

### Pharmacological inhibition of EXO1 displays anti-tumoral activity *in vivo*

To further explore EXO1 inhibition as a potential therapeutic strategy for EwS therapy, first we carried out clonogenic survival assay by treating A-673, RD-ES, and U2OS cells with C73. Notably, EwS cells were significantly more sensitive to C73 than U2-OS (Fig. 5A). Accordingly, treatment with C73 strongly increased the levels of the apoptotic markers cleaved PARP1 and cleaved caspase-3 in EwS cells compared to U2-OS (Fig. 5B).

**Figure 5.**
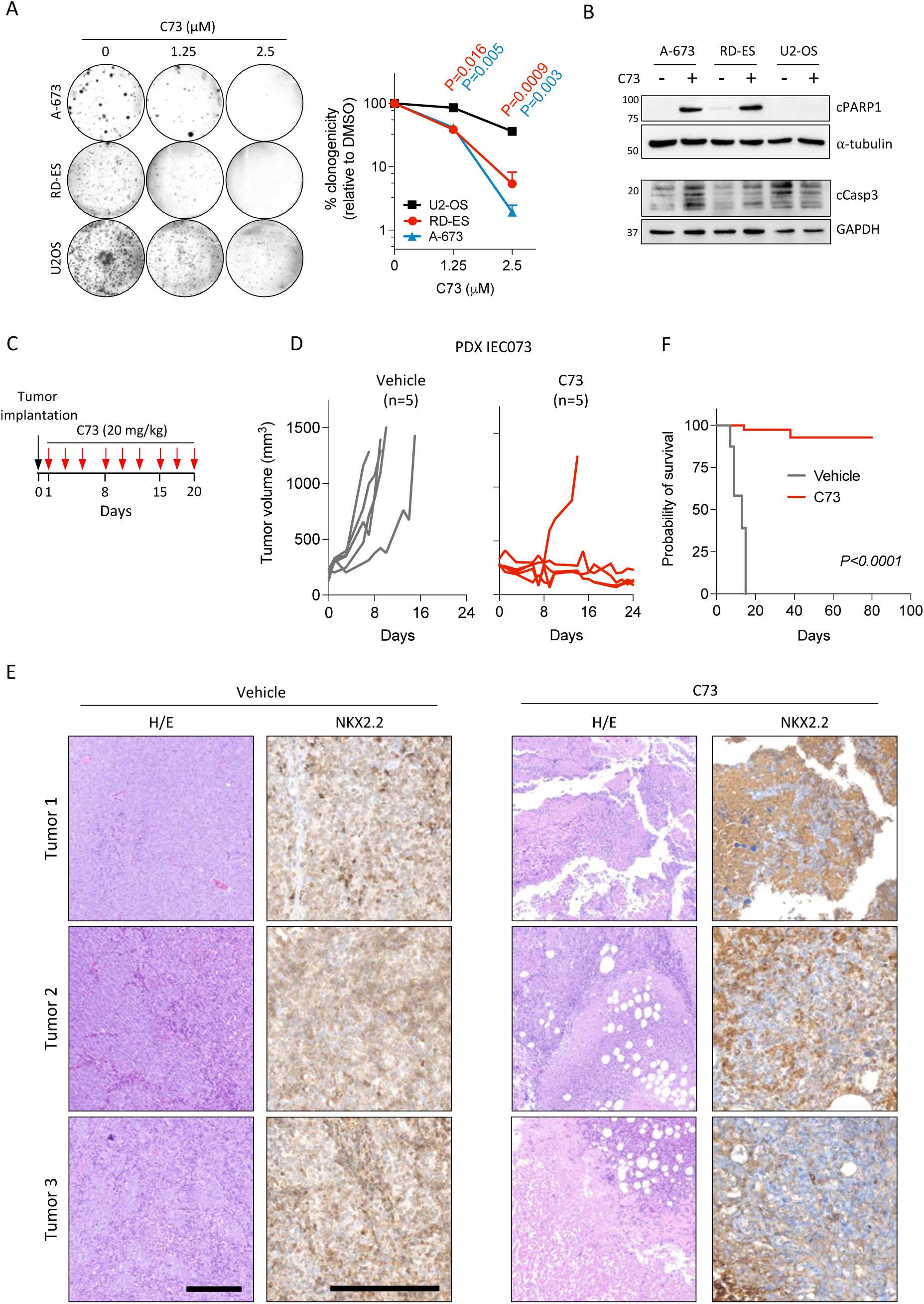
Pharmacological inhibition of EXO1 displays anti-tumoral activity *in vivo.* **(A)** Analysis of the effect of EXO1 inhibition on the clonogenic capacity of A-673, RD-ES, and U2-OS cell lines. Cells were incubated with indicated concentrations of C73 for 10-15 days. *Left*, representative images. *Right,* data represent the mean (+SEM) number of colonies (relative to DMSO); n≥3 independent experiments. *P* value was determined by two-tailed paired t-test. **(B)** Study of the induction of apoptotic markers, cleaved PARP1 and Caspase-3, after EXO1 inhibition by WB. When indicated, cells were incubated with 2.5 µM of C73 for 96 h. Molecular weight in kDa. GAPDH and α-tubulin: loading controls. **(C)** Scheme of the experimental settings of EXO1 inhibitor treatment *in vivo*. **(D)** Evaluation of tumor growth of PDX model IEC-073 over 23 days after the beginning of C73 treatment. **(E)** Immunohistological analysis of tumours after 12 days of the beginning of treatment. Representative images. Scale bar: 250 µm. **(F)** Kaplan-Meier survival analysis. *P* value was determined by the Mantel-Cox test.

Next, we explored the antitumoral activity of C73 *in vivo* using a EwS patient-derived xenograft model. Tumours were subcutaneously implanted and treated with doses of C73 that did not cause detectable weight loss, suggesting that C73 dosing was safe (Supplementary Fig. 5). Injections were administered in three-day intervals to reduce the vehicle (DMSO) toxicity (see methods for details) (Fig. 5C). Remarkably, treatment very significantly reduced tumor growth compared to vehicle, remaining almost undetectable (complete response) in 4 of 5 tumours (Fig. 5D). In accordance, immunohistological analysis of tumours after 12 days of the beginning of treatment revealed a very drastic decrease of cellularity (Fig. 5E). Notably, the remaining cells in C73-treated tumours no longer displayed the typical EwS cell morphology and were generally negative for NKX2.2, a well-described EwS marker^44^ (Fig. 5E). Importantly, 60 days after end of treatment 75% of tumours showing a complete response remained undetectable very significantly extending survival of host mice (Fig. 5F). Altogether, our findings highlight EXO1 inhibition as a promising therapeutic strategy for the treatment of EwS.

## DISCUSSION

Multimodal treatment approaches have substantially improved survival in EwS, particularly for patients with localized disease. However, long-term side effects of intensive cytotoxic regimens, along with poor outcomes in relapsed or metastatic disease, highlight the urgent need for novel therapeutic strategies^45^. Here, we explored an alternative approach based on EWSR1::FLI1 promoting replication stress and genome instability. We hypothesized that EwS cells would rely on DDR factors that mitigate the cytotoxic effects induced by the chimeric oncoprotein. Importantly, our results prove that EWS::FLI1 creates a dependence on EXO1.

An effect of EXO1 depletion on cell survival has been proven widely variable across other cancer cell lines, depending on their lineage or the presence of specific mutations. For instance, EXO1 is essential for the survival of BRCA1-deficient cells^34,46^. Additionally, *EXO1* silencing impairs the clonogenic growth of hepatic or prostate cancer cells^47,48^, apparently, in the absence of mutations affecting other factors involved in homologous recombination repair. Nevertheless, this effect has not been formally attributed to the induction of cell death, but rather to impaired cellular proliferation^47,48^. We demonstrate that *EXO1* silencing in EwS cells triggers cell death, impairing clonogenicity and tumor growth. Moreover, our results show that, despite variability in *EXO1* levels among the employed EwS cells, they are similarly affected by EXO1 depletion or inhibition, accentuating the strong EwS dependency on EXO1. Strikingly, our findings are inconsistent with public data from genome-wide CRISPR-Cas9 screens, which show a slightly higher, although not significant dependency of EwS cells on EXO1, in comparison with other tumoral cell lines (data from DepMap portal, CRISPR (Chronos)).

Our results prove that EXO1 depletion and inhibition induces DSBs in EwS, in contrast to other normal and tumoral cells in which EXO1 loss does not significantly increase the levels of DNA damage in the absence of exogenous genotoxic agents^35,49–52^. EXO1-mediated long-range resection of DNA ends is a key step of homologous recombination pathway^26^. Moreover, it has been recently shown that, in the absence of BRCA1, EXO1 is essential for SSA repair^34^. However, EXO1 depletion in EwS cells did not accumulate DSBs in S but in G1-phase. Given that Kooij *et al.* showed that in the absence of BRCA1, EXO1 loss induces DNA lesions in S-phase, these results suggest that it is unlikely that EwS dependency on EXO1 might be caused by the previously reported ‘BRCAness’^22^. Importantly, our results also demonstrated the presence of DNA lesions in mitotic chromosomes. For the last years, numerous reports have illustrated that EWSR1::FLI1 promotes replication stress^20–22^. Importantly, replication stress caused by human oncogenes slows replication forks, leading to UR-DNA in mitosis, which becomes a known source of DNA damage that triggers MiDAS repair^53–55^. Accordingly, our findings show that EWSR1::FLI1 leads to the accumulation of late replication intermediates in mitosis and activates a RAD52-dependent MiDAS. The functions of EXO1 during mitosis have been poorly addressed. More importantly, the potential role of EXO1 in MiDAS repair remained unknown. In response to DNA damaging agents, EXO1 is degraded in G2/M phases to prevent DNA end resection^56^. Nonetheless, our findings indicate that EXO1 plays a role in DNA repair during cell division, in the absence of exogenous sources of DNA lesions, preventing EWSR1::FLI1-associated cytotoxicity and mitotic genome instability. Given that MiDAS is a BIR-mediated process, we hypothesized EXO1 function in mitotic DNA synthesis might be similar to its activity during BIR repair. Interestingly, two different reports have shown that Exo1 inhibits BIR in *S. cerevisiae*^57,58^. However, in the absence of Tel1 (the yeast homologous of ATM, the ataxia-telangiectasia mutated gene), Exo1 extends 3’-ended ssDNA facilitating the search of homologous sequences. This model is particularly interesting considering that, in cells lacking *TEL1*, MRX complex generates an Exo1 substrate which is similar to the intermediate generated by MUS81-dependent cleavage during MiDAS^53^. Notably, our analysis of UFBs indicated that EXO1 loss does not affect the processing of late replication intermediates, but rather impairs the formation of recombination intermediates, as indicated by the reduction of RPA-coated UFBs. Thus, we reason that, upon MUS81-cleavage, EXO1-dependent extensive resection of single-ended DSBs, may favor 3’-ended strand invasion and subsequently, DNA synthesis. While other nucleases, such as WRN, have recently been implicated in this role, it remains unclear whether EXO1 activity is redundant or uniquely restricted to EwS^59^.

From a translational perspective, our findings are especially relevant. EXO1 inhibition with C73 phenocopied the effects of genetic depletion in EwS cells and showed marked antitumoral activity *in vivo*, including profound responses in a patient-derived xenograft model. These results provide proof of principle that EXO1 is pharmacologically actionable in EwS. At the same time, the translational implications of these experiments should be interpreted with appropriate caution. C73 represents an important first-in-class tool compound, but further work will be needed to define its selectivity, pharmacokinetic and pharmacodynamic properties, therapeutic window, and possible off-target effects. The growing interest in, and the increased specificity and solubility of EXO1 inhibitors beyond C73 will facilitate the necessary assays to advance studies evaluating EXO1 as a therapeutic target in EWS in the near future^60^. Broader testing across additional EwS models will also be necessary to establish the robustness and generalizability of the response.

Our study also raises the possibility that EXO1 may have clinical utility beyond its role as a therapeutic target. Prior studies have revealed an association between elevated *EXO1* expression and a higher risk of patient’s death, metastasis or reduced immune infiltration, in other tumors^61–63^. Remarkably, we found that high EXO1 levels correlate with poor patient outcome and is associated with chr1q gain, a genomic feature previously validated as a marker of poor prognosis in EwS. This suggests that EXO1 may reflect, at least in part, a more aggressive and genomically stressed disease state. Whether EXO1 expression, chr1q status, EWSR1::FLI1 activity, or functional markers of replication stress and MiDAS could serve as predictive biomarkers of response to EXO1-directed therapies remains to be determined. This will be an important question for future translational studies.

Finally, our findings open the door to rational combination strategies. Because EXO1 appears to support tolerance to fusion-driven replication stress, its inhibition may be particularly effective when combined with agents that further increase replication stress or limit compensatory stress-response pathways. In this regard, combinations with ATR inhibitors are particularly attractive, given their established activity in EwS^20^ and their potential to increase dependence on mitotic rescue pathways such as MiDAS^42^. Likewise, targeting complementary components of MiDAS may further enhance the therapeutic effect of EXO1 inhibition.

In summary, our study identifies EXO1 as a non-oncogene dependency created by EWSR1::FLI1 in EwS. We show that EXO1 protects EwS cells from mitotic genome instability, likely by facilitating efficient MiDAS-associated processing of fusion-induced late replication intermediates, and we provide preclinical evidence that EXO1 inhibition has therapeutic potential in this disease. Together, these findings nominate the EXO1/MiDAS axis as a mechanistically grounded and translationally relevant vulnerability in EwS.

## METHODS

### Cell lines and culture conditions

Cell lines characteristics, origin and culture conditions are depicted in Supplementary Table 2. Growth media was supplemented with 10% fetal bovine serum (FBS, Thermo Fisher, 10270106) and 1% penicillin/streptomycin (Thermo Fisher, 15140122). A-673/TR/shEF cell line was gently provided by J. Alonso (Madrid, Spain)^38^. HeLa Tet-On EF (HeLa EF) model was previously generated by our group^30^. Cell lines were grown at 37°C, 5% CO_2_, routinely tested for mycoplasma contamination using the MycoAlert Detection Kit (Lonza Group Ltd, LT07-318) and authenticated by STR analysis (CLS, Cell Lines Service GmbH). All commercial cell lines were obtained from ATCC, and non-commercial, from HHU Düsseldorf (EuroBoNet Project). A-673/TR/shEF and HeLa EF cells were incubated with 1 µg/ml of doxycycline (DOX, Sigma-Aldrich, D9891) for indicated times. Drugs used in this study are listed in Supplementary Table 2.

### Generation of DOX-inducible shRNA-expressing cellular models

Non-targeting negative control and a specific shRNA targeting EXO1 were cloned into the lentiviral pLKO-Tet-on-all-in-one vector system (Addgene, 21915), according to available protocol^64^. Oligonucleotide sequences are shown in Supplementary Table 2. Lentiviral particles were produced in HEK293T cells. RD-ES cells were infected with HEK293T supernatants and selected with 1.5 µg/ml puromycin (InvivoGen). Knockdown efficacy was confirmed by qPCR and WB after 72 h of DOX incubation (1 µg/ml).

### RNA extraction, reverse transcription and qPCR

RNA was extracted with the miRNeasy Kit (Qiagen, 217084), according to the manufacturer’s instructions. For DNA removal, columns were incubated with RQ1 DNAse (Promega, M6101). RNA was retrotranscribed using the High Capacity cDNA Reverse Transcription Kit (Applied Biosystems, 4368814). qPCR was performed using the iTaq Universal SYBR Green Supermix (Bio-Rad, 1725121) and TaqMan probes shown in Supplementary Table 2. qPCR data were analyzed using ExpressionSuite software v1.3 (Thermo Fisher). 2^-ΔCt^ method was used to calculate relative gene expression levels, where ΔCt correspond to the difference between the Ct of the target gene and the Ct of *GAPDH*.

### Transient transfection

For siRNAs transfection (listed in Supplementary Table 2), cells were seeded in 24-well or 6-well plates at a density of 2-5 x 10^4^ or 1.5-2.5 x 10^5^ per well, respectively. After 24 h, cells were transfected with indicated siRNAs (20-40nM) using jetPRIME (Polyplus, 101000046), according to the manufacturer’s protocol. Cells were cultured for 48-72 h prior experiments. Knockdown efficacy was confirmed by qPCR.

### Colony formation assay

Cells were seeded in 6-well plates at a density of 1-3 x 10^3^ cells per well. Then, DOX (1µg/ml) or C73 (indicated concentrations) were immediately added to the medium. For clonogenic assays using siRNAs, cells were transfected in 6-well plates as previously indicated. After 48-72 h of transfection, cells were detached, counted, and seeded in 6-well plates at a density of 1-3 x 10^3^ cells per well in fresh medium. After 7-15 days, cells were washed with PBS, and fixed/stained with 2% methylene blue in 70% ethanol for 30 min. Cell plates were washed with distilled water and colonies were counted manually.

### Sphere formation assay

For the analysis of anchorage-independent growth, cells were seeded in 96-well ultra-low attachment microplates (Corning, 7007) at a density of 2 x 10^3^ cells per well (3 wells per condition). DOX (1µg/ml) was immediately added to the medium. After 6 days, tumorspheres were photographed using Olympus IX-71 inverted microscope. The area of tumorspheres was determined using ImageJ software version 1.54.

### Metaphase spreads

For the preparation of metaphase spreads, cells were seeded in 100 mm dishes at low confluence. After 24 h, demecolcine solution (Sigma-Aldrich, D1925) was directly added to the culture medium at a final concentration of 0.2 μg/ml. Cells were grown in the presence of demecolcine for 6-14 h (depending on the proliferation rate). Then, cells were collected and incubated in a 0.03 M trisodium citrate solution at 37 °C for 30 min. After that, cells were fixed with 3:1 methanol-acetic acid. The fixation solution was added dropwise under rotation and, subsequently, cells were centrifuged at 750 x g for 5 min at 4 °C. Fixation step was repeated 3 times. Finally, fixed cells were dropped onto acetic acid-humidified slides. For the analysis of chromosome breaks, slides were stained with 5% Giemsa solution (Sigma-Aldrich, GS500) for 20 min, and mounted using DPX (Sigma-Aldrich, 06522).

### Fluorescence in situ hybridization (FISH)

For whole chromosome FISH, metaphase spreads were generated as previously indicated. For chr1q FISH, cells were directly fixed in 3:1 methanol-acetic acid, and dropped onto acetic acid-humidified slides. FISH probes are shown in Supplementary Table 2. Slides were immersed in 2x SSC for 2 min at room temperature and then, dehydrated in an ascending ethanol series (70%, 85% and 96%) for 2 min each, at room temperature. Probes were added onto the preparation and denatured for 2 min at 75°C. After an overnight incubation at 37 °C, slides were immersed 5 min in 2x SSC at room temperature, 2 min in 0.4x SSC at 72 °C and 30 sec in 2x SSC at room temperature. Finally, slides were counterstained with DAPI (Sigma-Aldrich, 09542) for 5 min and mounted using Dako Fluorescent Mounting Medium (Dako). Slides were visualized using Olympus BX61 Fluorescence Motorized Microscope.

### Analysis of cell cycle and apoptosis by FACS

For the analysis of cell cycle, cells were seeded at a density of 0.5-1 x 10^6^ cells in 100 mm dishes. 10 µM EdU (BaseClick, BCN-001) was added to the culture medium 20 min before the end of DOX incubation. Then, cells were washed twice with PBS, detached and fixed with 4% PFA for 10 min at room temperature. After fixation, cells were permeabilized with PBS-0.2% Triton X-100 for 10 min. After washout, cells were incubated with the Reaction cocktail solution (100 mM Tris-HCl pH 8.5, 1 mM CuSO_4_, 1 µM Fluorescein Azide 6-FAM (BaseClick, BCFA-001), 100 mM ascorbic acid) for 30 min at room temperature, under agitation and in the dark. Then, cells were washed twice with PBS-1% BSA and 4 times with PBS-0.1% Tween 20 for 10 min. Finally, cells were incubated with propidium iodide (PI) solution (100 µg/ml PI and 100 µg/ml RNAse A) for 30 min. For the analysis of apoptosis, cells were seeded at a density of 0.5-1 x 10^6^ cells in 100 mm dishes. After 96 h of DOX incubation, cells and supernatants were collected and incubated with Annexin V and PI (Immunostep, FITC-conjugated, ANXVKF7-100T) according to the manufacturer’s instructions for 15 min, in the dark. 10,000-20,000 cells were inspected for each sample using a BD FACSCanto II flow cytometer. Data were analyzed using FlowJo v.10.2 software. Total Annexin V-positive cells (early + late apoptosis) were shown.

### Analysis of DNA damage by FACS

For the analysis of DNA damage in S-phase population, cells were seeded at a density of 0.5-1 x 10^6^ cells in 100 mm dishes. 10 µM EdU was added to the culture medium 20 min before the end of DOX incubation. Fixation, permeabilization, blockage and click chemistry reaction was performed as previously indicated. Then, cells were incubated with anti-!H2AX antibody (Millipore, 05-636), diluted 1:1000 in PBS-1% BSA for 2 h, under rotation; and subsequently, with anti-mouse Alexa Fluor 488-conjugated secondary antibody (Invitrogen, A11001) for 1 h. For the analysis of DNA damage in mitosis, cells were incubated with anti-!H2AX and anti-pH3(S10)(Cell signaling, 3377) antibodies diluted 1:1000 in PBS-1% BSA for 2 h; and then, with anti-mouse Alexa Fluor 488-conjugated and anti-rabbit Alexa Fluor 546-conjugated secondary antibodies (Invitrogen, A11035) for 1 h. Finally, cells were inspected using a BD FACSCanto II flow cytometer. Analysis of FACS data was performed using FlowJo v.10.2 software.

### Immunofluorescence (IF)

IF assay was performed as previously described by our group^65^. 2-5 x 10^4^ cells were seeded on coverslips in 24-well plates. For EwS cell lines, coverslips were pre-incubated with 10 µg/ml human fibronectin (Corning, 356008) overnight at 4°C. Briefly, cells were fixed with 4% PFA for 10 min at room temperature, permeabilized with PBS-0.2% Triton X-100 for 10 min, and blocked with PBS-5% BSA for 1 h. Then, cells were incubated with primary antibodies (shown in Supplementary Table 2) for 1-3 h in PBS-1% BSA. After that, cells were incubated with corresponding Cy2/Cy3-conjugated secondary antibodies (Jackson Immunoresearch) for 1 h, diluted 1:1000 in PBS-1% BSA. After washes, cells were counterstained with DAPI for 5 min and mounted using Dako Fluorescent Mounting Medium (Dako). Images were acquired using Olympus BX61 Fluorescence Motorized Microscope and analyzed with ImageJ software version 1.54. To determine !H2AX nuclear intensity, a mask was generated using DAPI and the “analyze particles” tool. Then, mask was applied to !H2AX images. The intensity of each nuclei was divided to the nuclear area.

### Western blot (WB)

WB was performed as previously described by our group^66^. Briefly, cell lines were lysed using RIPA buffer supplemented with protease (Roche, 11697498001) and phosphatase inhibitors (1 mM Na_3_O_4_V (Sigma-Aldrich, 450243)) and 1 mM NaF (Sigma-Aldrich, S1504)). Primary antibodies (listed in Supplementary Table 2) were blocked in Tris buffered saline buffer 0.1% Tween-20; 5% BSA. HPRT-conjugated anti-mouse (Biorad, 1706516) and anti-rabbit (Biorad, 1706515) secondary antibodies were incubated for 1 h at room temperature (1:5000). Images were acquired in Chemidoc Imaging System (Bio-Rad) using Pierce ECL western blotting substrate (Thermo Fisher Scientific). Band density was determined using Image Lab Software (Bio-Rad).

### Tissue microarrays and immunohistochemistry (IHC)

IHC was performed at HUVR-IBiS Biobank facility according to conventional protocols, previously described by our group^65^. After blockage, sections were incubated with anti-EXO1 (Affinity Biosciences, DF3615) at 1:200 or anti-NKX2.2 (Sigma-Aldrich, EP336); and with peroxidase-labeled secondary antibodies and 3,3-diaminobenzimide, according to manufacturer’s protocol (Leica Biosystems). Slides were counterstained with Hematoxylin and mounted using DPX (BDH Laboratories). Images were acquired using Olympus BX61 Fluorescence Motorized Microscope.

For the evaluation of the association between EXO1 levels and EwS overall survival, EXO1 IHC was performed in a TMA comprising 76 clinically annotated FFPE EwS tumors. All identified patients and collected data were in accordance with guidelines of MDACC and IBIS’s institutional review board. EXO1 IHC was evaluated by an experienced pathologist. The staining score (S, 0 to 300) was determined as the product of the EXO1 signal intensity (0 to 3) and extension (0 to 100%). Samples were grouped into: EXO1 high (S ≥ 250, n=25) and, medium (S > 100 and < 250), low and negative (S ≤ 100)(n=51). *P* value was determined by Mantel-Cox test using GraphPad PRISM version 9 (GraphPad Software Inc., CA, USA).

### Evaluation of the association between DDR factors and EwS overall survival

Survival analysis was performed as previously described^24^, in a cohort of 166 EwS transcriptomes (GSE63157, GSE34620, GSE12102, GSE17618) with available clinical annotations. Samples were divided into tertiles according to the expression levels of the corresponding gene. *P* value was determined by Mantel-Cox test using GraphPad PRISM version 9 (GraphPad Software Inc., CA, USA) and adjusted by Bonferroni correction.

### Analysis of published microarray expression data

Public microarray expression data were obtained from Gene Expression Omnibus (GEO, NCBI). Accession numbers are shown in Supplementary Table 1. Gene expression analysis was performed using R (version 4.2.2) and “affy” package from Bioconductor. Expression data were normalized using the robust multi-array average (RMA) method.

### Generation of chr1q gain expression signature

To generate an EwS-specific chr1q-gain expression signature we performed a differential expression analysis between chr1q gain (CHLA-10, EW-22, EW-24, EW-7, MHH-ES-1, SK-ES-1, TC-32) and chr1q normal (A-673, EW-1, POE, RH1, SK-N-MC) EwS cell lines, using GEO2R tool from NCBI. Data were obtained from Ewing Sarcoma Cell Line Atlas (ESCLA)(GSE176190)^67^. Then, we selected genes located in chr1q and significantly overexpressed in chr1q gain with respect to chr1q normal cells (Log2 FC > 1, *P* value < 0.05). 166 clinically annotated EwS transcriptomes (GSE63157, GSE34620, GSE12102, GSE17618) were scored according to chr1q-gain expression signature using ssGSEAProjection (v4)(GenePattern)^68^. Samples were grouped into “high” (n = 30, 18% of samples, according to the expected percentage of chr1q gain) and “low” chr1q gain (n = 136).

### RD-ES/TR/shEXO1 xenograft model

RD-ES/TR/shEXO1 cell suspension was mixed 1:1 with Corning Matrigel Basement Membrane Matrix (Sigma-Aldrich, CLS354234). 5 x 10^6^ cells were subcutaneously injected in the flank of nu/nu athymic nude mice (Envigo). When tumors reached 1500 mm^3^, mice were sacrificed by cervical dislocation, tumors were excised, and pieces of around 5×5 mm were reimplanted. When tumor volume reached ∼150 mm^3^, mice were randomized into two groups. DOX was administered for 20 days via drinking water at 2 mg/ml + 5% sucrose. The control group received only sucrose. Drinking water was replaced every 3 days. Tumor size (using an electronic caliper) and mice weight were measured every two days. Tumor volume was calculated as *V* = (*a* x *b*^2^ x π)/6, where *b* is the smallest measure and *a* the largest measured.

### C73 inhibitor treatment in vivo

To evaluated the antitumoral activity of C73, we employed the IEC-073 EwS PDX model. Tumors were excised from mice and cut into approximately 5×5 mm pieces. Then, tumors were subcutaneously implanted in nu/nu athymic nude mice (Envigo). When tumor volume was ∼150 mm^3^, mice were randomized into two groups and treated with C73 (20 mg/kg/day) intraperitoneally, every two days for three weeks. Control group was treated with physiological saline solution. Mice were sacrificed by cervical dislocation once tumors reached 1200-1500 mm^3^, for ethical reasons. Some animals were sacrificed on day 12 of treatment and tumors were embedded in paraffin for the evaluation of tumoral necrosis. Hematoxylin/Eosin (H/E) staining was performed according to conventional protocols and evaluated by an experienced pathologist. NKX2.2 IHC was performed as previously indicated. Images were acquired using Olympus BX61 Fluorescence Motorized Microscope. Survival curves were generated using GraphPad Prism software v.9. (GraphPad, San Diego, CA, USA), and *P* value was determined by Mantel-Cox test. Animal experiments were approved by the Consejeria de Agricultura, Pesca, Agua y Desarrollo Rural; Junta de Andalucia (08-05-2018-082) and performed according to protocols and conditions reflected in the European guidelines (EU Directive 2010/63/EU).

### Statistical analysis

Statistical analyses were performed using GraphPad PRISM version 9 (GraphPad Software Inc., CA, USA). *P* values are indicated in the figures, and the statistical test used for comparisons are specified in the corresponding figure legends. *In vitro* experiments were conducted using three or more independent replicates. For *in vivo* studies, the number of animals was minimized while ensuring robust and reliable results, in accordance with the principles of the Three Rs (Replacement, Reduction, and Refinement).

## CONFLICT OF INTEREST

Authors declare no conflicts of interest.

## ACKNOWLEDGEMENTS

We thank to the HUVR-IBiS Biobank facility for IHC support.

## AUTHOR CONTRIBUTION

J.O-P. performed most of the experiments unless indicated, analyzed data and generated the figures. L.L-S. and D.D-B. helped with *in vitro* experiments. C.J-P., P.G-P. and M.P. supported animal experiments. F.H.G. and M.J.C-G helped with the generation of RD-ES/TR/shEXO1 cellular model. J.A. provided A673/TR/shEF cellular model. L.Z. and B.S. provided C73 inhibitor. T.G.P.G. provided EwS patients microarray expression data and clinical information, provided laboratory infrastructure and financial support for J.O-P. international internship and biological guidance. E.dA., F.G-H. and J.O-P. designed experiments and wrote the manuscript. E.dA. and F.G-H. coordinated and supervised the study. All authors reviewed the manuscript.

## FUNDING

The laboratory of E.dA. is supported by grants from ISCIII-FEDER (PI20/00003, PI23/1460, and PMP22/00054), Consejería de Salud y Consumo, Junta de Andalucía (PE-0186-2018, PI-0061-2020), Fundación Científica AECC (ECAEC222952DEAL), Fundación CRIS Contra el Cáncer, Asociación Pablo Ugarte, Fundación María García Estrada, and CIBERONC. The laboratory of F.G-H is supported by grants funded by MCIN/AEI/10.13039/501100011033/FEDER, UE (R+D+i PID2019-105212GB-I00 and PID2022-137143NB-I00) and the Andalusian Regional Government: Fondo Europeo de Desarrollo Regional (FEDER) y Consejería de Transformación Económica, Industria, Conocimiento y Universidades de la Junta de Andalucía (P20_00561) to F.G-H. J.O-P is supported by Ph.D. grant *Plan Propio* from the University of Seville. D. D-B. is supported by *Consejería de Economía, Innovación, Ciencia y Empleo, Junta de Andalucía* (PAIDI 2020, POSTDOC_21_00865). The laboratory of JA is supported by Instituto de Salud Carlos III (grant numbers PI20CIII/00020, DTS22CIII/00003, PMP21-00073); Asociación Pablo Ugarte (TRPV205/18, DGDO195722); Asociación Candela Riera, Asociación Todos Somos Iván & Fundación Sonrisa de Alex (TVP333-19, TVP-1324/15). The laboratory of T.G.P.G. is supported by grants from the Matthias-Lackas Foundation, the Dr. Leopold und Carmen Ellinger Foundation, the Deutsche Forschungsgemeinschaft (DFG 458891500), the German Cancer Aid (DKH-7011411, DKH-70114278), the Boehringer-Ingelheim foundation, the Dr. Rolf M. Schwiete foundation, the SMARCB1 association, the Ministry of Education and Research (BMBF; SMART-CARE and HEROES-AYA), and the Barbara and Wilfried Mohr foundation.

## FIGURES

**Supplementary Figure 1.**
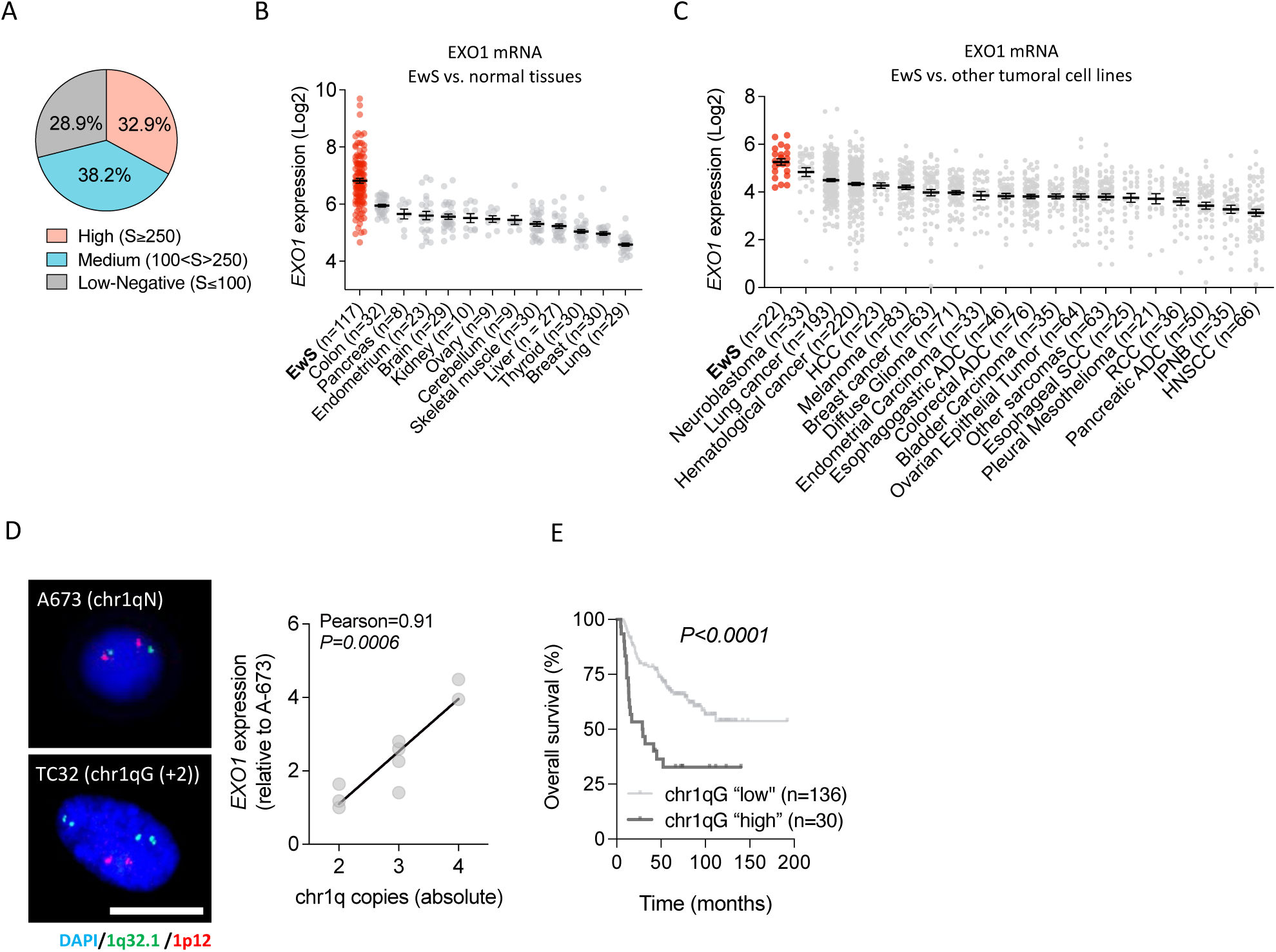
(**A**) Distribution of 76 EwS samples according to EXO1 staining score (S)(intensity x extension). **(B)** *EXO1* expression levels (mean ± SEM) in EwS samples and normal tissues. **(C)** Similar to (B) in EwS cells and other tumoral cell lines. Data were obtained from *DepMap, Broad (2025). DepMap Public 25Q3. Dataset (depmap.org*). **(D)** Evaluation of the correlation between *EXO1* expression levels (determined by qPCR) and chr1q copy number in 9 EwS cell lines. *Left,* representative FISH images. Green: chr1q32.1 (*MDM4* gene), red: chr1p12. DAPI counterstain. Scale bar: 10 µm. *Right,* correlation plot. Linear correlation was determined by Pearson’s correlation coefficient. **(E)** Kaplan-Meier survival analysis of 166 EwS samples stratified by chr1q expression signature. *P* value was determined by Mantel-Cox test.

**Supplementary Figure 2.**
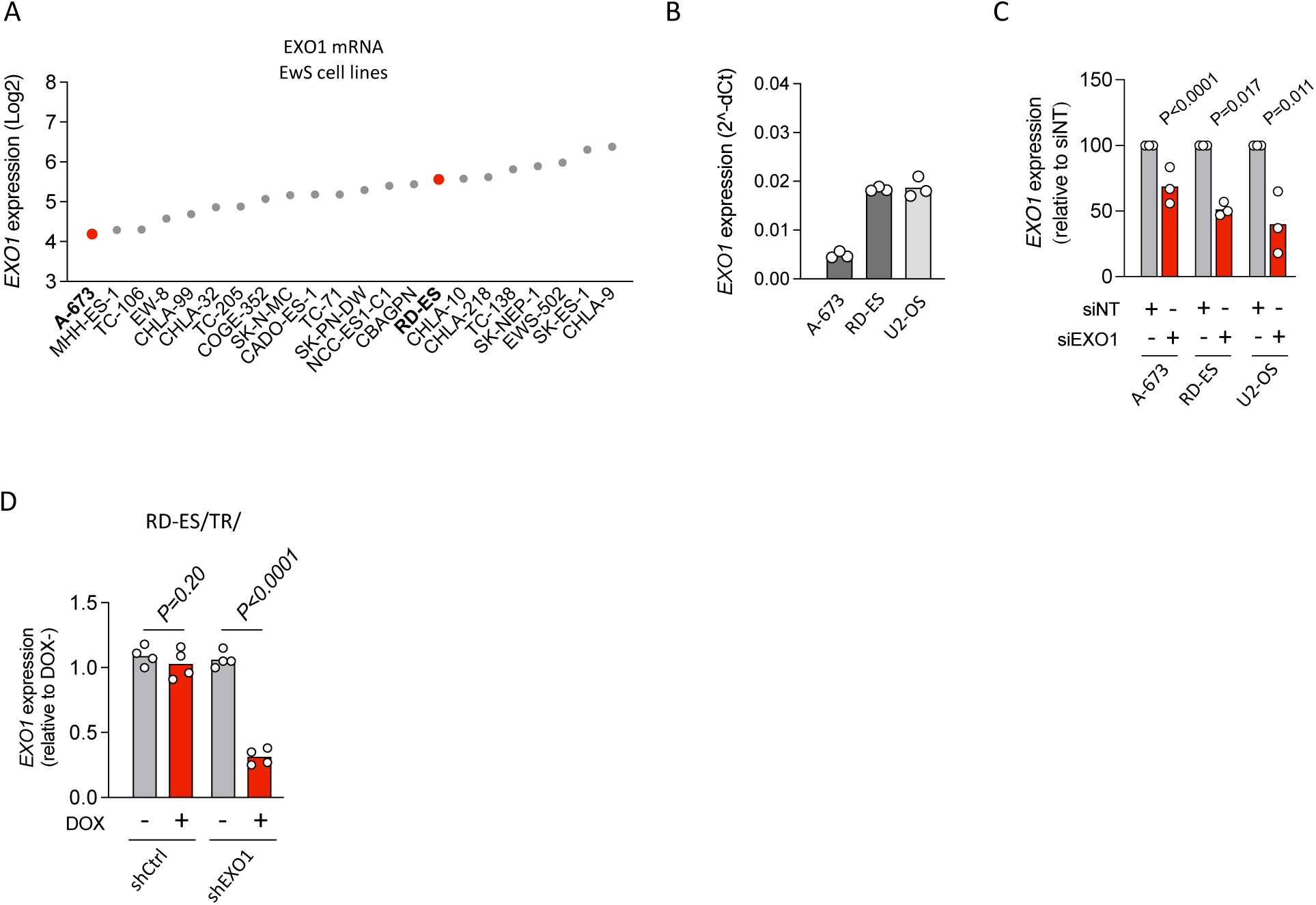
*EXO1* expression levels in EwS cell lines. Data were obtained from *DepMap, Broad (2025). DepMap Public 25Q3. Dataset (depmap.org)*. **(B)** *EXO1* expression levels in A-673, RD-ES, and U2-OS cells, determined by qPCR (normalized to *GAPDH*). Data are the mean of 3 independent experiments. **(C)** Evaluation of *EXO1* knockdown in A-673, RD-ES and U2-OS after 72 h of siRNA transfection by qPCR. Data are the mean of *EXO1* expression levels (normalized to *GAPDH* and relative to siNT); n=3 independent experiments. **(D)** Evaluation of EXO1 expression levels in RD-ES/TR/shEXO1 (and shCtrl) cells after 72 h of DOX incubation by qPCR. Data represent the mean of *EXO1* expression (normalized to *GAPDH* and relative to DOX-); n=4 independent experiments. *P* value was determined by a two-tailed paired t-test.

**Supplementary Figure 3.**
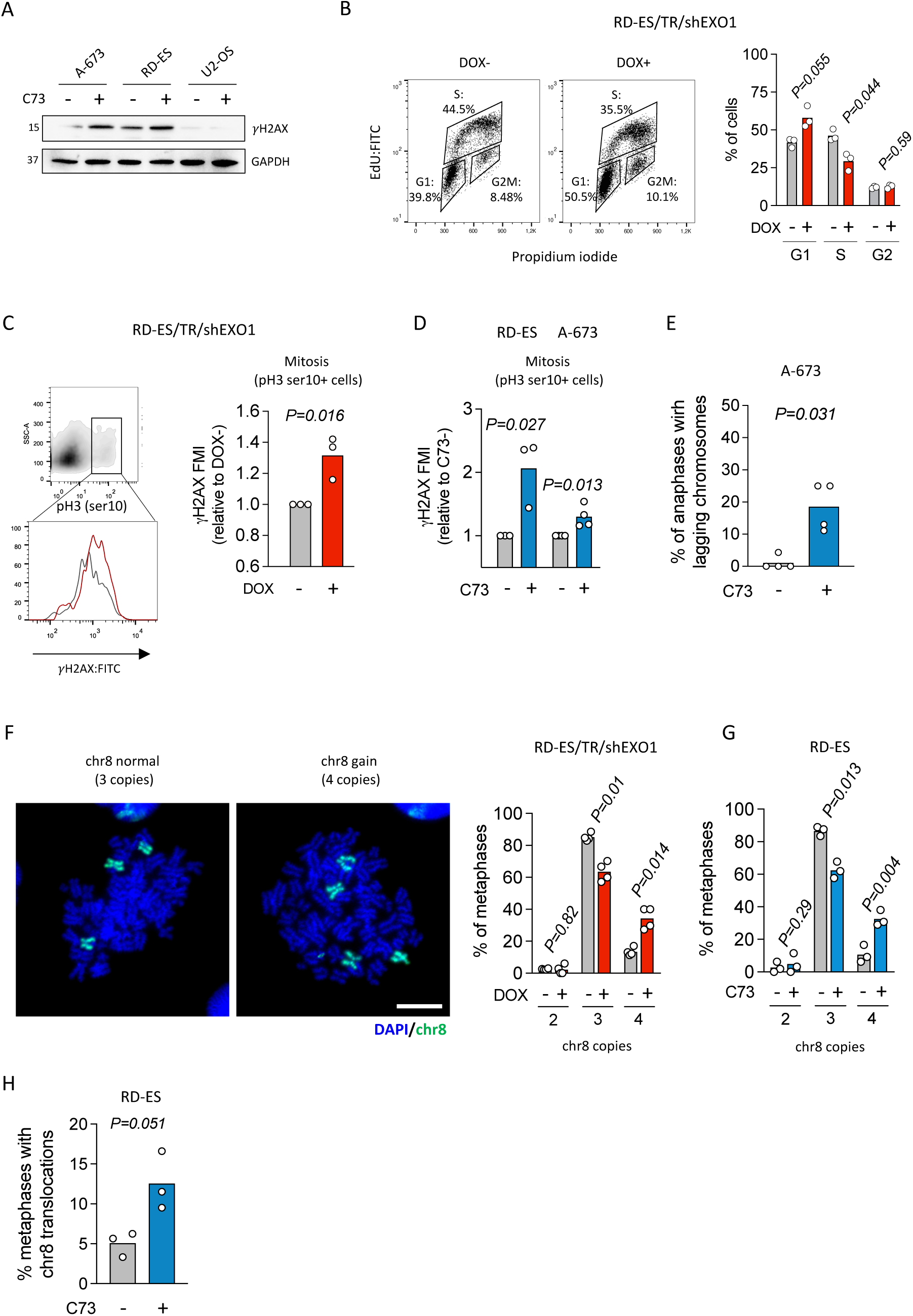
(**A**) Analysis of the effect of EXO1 inhibition on the induction of DSBs by !H2AX WB. When indicated, cells were treated with 2.5 µM of C73 for 72 h. Molecular weight in kDa. GAPDH: loading control. **(B)** Analysis of the effect of *EXO1* silencing on cell cycle distribution by EdU-PI FACS. When indicated, cells were incubated with DOX for 48 h. *Left,* representative dot plots. *Right*, data represent the mean percentage of cells per phase; n=3 independent experiments. **(C)** Evaluation of the induction of DSBs in mitosis upon EXO1 depletion by !H2AX FACS. pH3S10 was used as a mitotic marker. *Left,* histogram shows !H2AX intensity in pH3S10-positive (mitotic) cells. *Right,* data represent the mean of !H2AX FMI (relative to DOX-) in pH3S10-positive cells; n=3 independent experiments. **(D)** Similar to (C) after incubation with 2.5 µM of C73 for 72 h; n≥3 independent experiments. **(E)** Effect of EXO1 inhibition on the levels of lagging chromosomes. Cells were treated with 2.5 µM of C73 for 72 h. Data represent the mean of the frequency of anaphases showing lagging chromosomes; n=4 independent experiments. **(F)** Effect of EXO1 depletion on numerical chromosomal aberrations by chr8 FISH. Cells were incubated with DOX for 72 h. *Left*, representative images of metaphases with 3 (“normal”) or 4 (gain) copies of the chr8. DAPI counterstain. Scale bar: 50 µm. *Right*, data represent the mean of the percentage of metaphases with indicated copies of chr8. **(G)** Similar to (F) after treatment with 2.5 µM of C73 for 72 h; n=3 independent experiments. **(H)** Analysis of the effect of EXO1 inhibition on the formation of chromosomal translocations by chr8 FISH. Cells were incubated with 2.5 µM of C73 for 72 h; n=3 independent experiments. *P* value was determined by two-tailed unpaired t-test in panels C and D; and paired t-test in panels B, E, F, G and H.

**Supplementary Figure 4.**
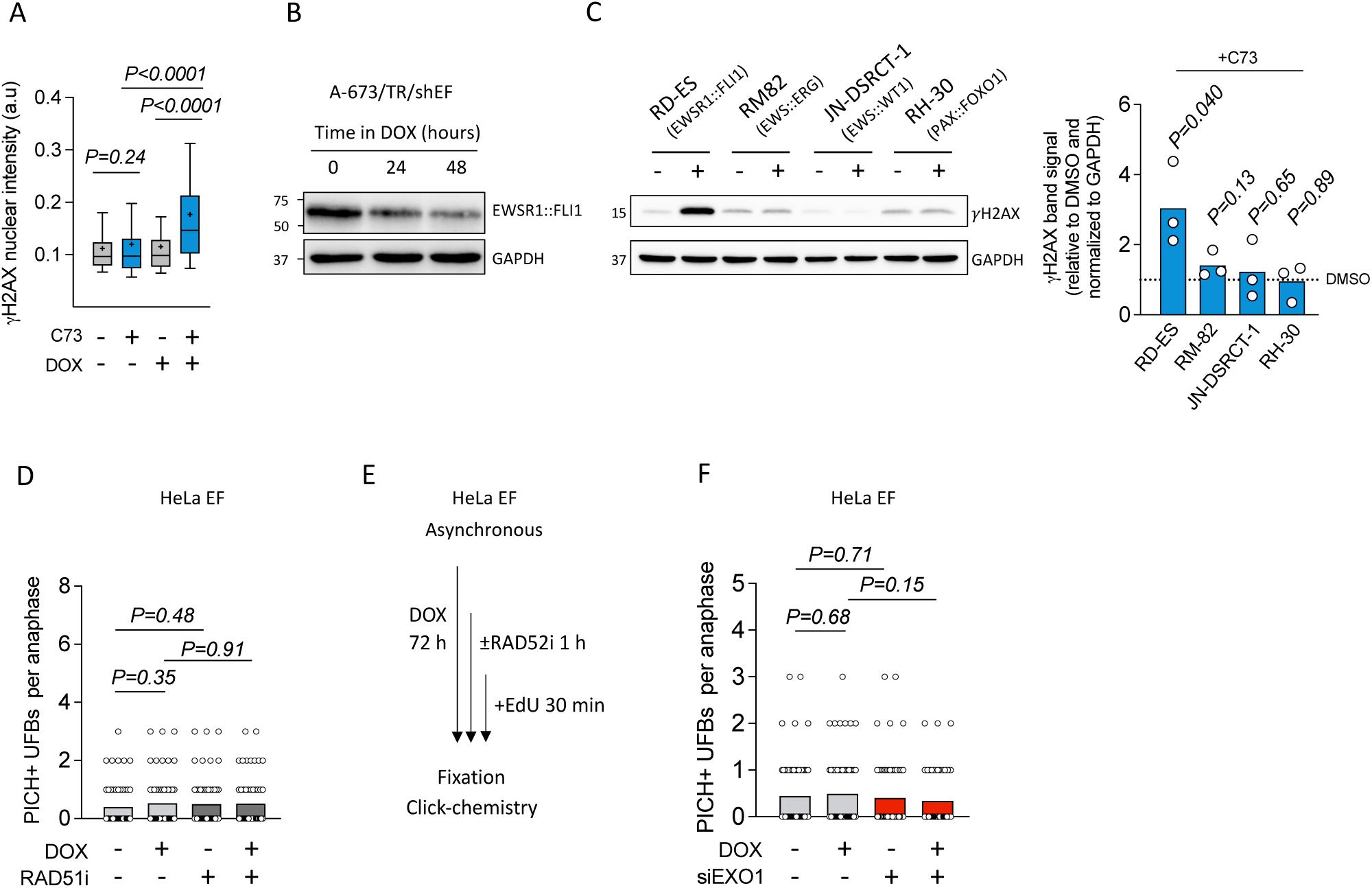
(**A**) Analysis of the relationship between EXO1 and EWSR1::FLI1 on the induction of DNA damage by !H2AX IF. When indicated, RD-ES/TR/shEXO1 cells were incubated with DOX for 72 h and treated with 2.5 µM of C73 for 48 h. Data represent the mean (+)(±10-90 percentile) of nuclear !H2AX intensity; n=3 independent experiments (∼140 cells were analyzed per replicate). **(B)** Evaluation of EWSR1::FLI1 protein levels in A-673/TR/shEF cells after incubation with DOX by WB. Molecular weight in kDa. GAPDH: loading control. **(C)** Evaluation of the induction of !H2AX after EXO1 inhibition by WB. When indicated, cells were treated with 2.5 µM of C73 for 72 h. *Left,* representative blots. Other details as in (B)*. Right*, data represent the mean of !H2AX band intensity (normalized to GAPDH) in C73-treated cells (relative to DMSO); n=3 independent experiments. **(D)** Evaluation of the effect of EWSR1::FLI1 ectopic expression on the levels of PICH-coated UFBs. When indicated, HeLa EF cells were incubated with DOX for 72 h and/or 10 µM of RAD51 inhibitor RI-1 for 1 h. Data represent the mean number of PICH-positive UFBs per anaphase; n=3 independent experiments (20 cells were analyzed per replicate). **(E)** Scheme of the protocol followed for the study of MIDAS efficiency. **(F)** Study of the effect of *EXO1* downregulation on EWSR1::FLI1-induced PICH-positive UFBs. When indicated, HeLa EF cells were incubated with DOX for 72 h and transfected with siNT or siEXO1 for 48 h. Data represent the mean number of PICH-positive UFBs per anaphase; n=3 independent experiments (∼25 cells were analyzed per replicate). *P* value was determined by two-tailed unpaired t-test.

**Supplementary Figure 5.**
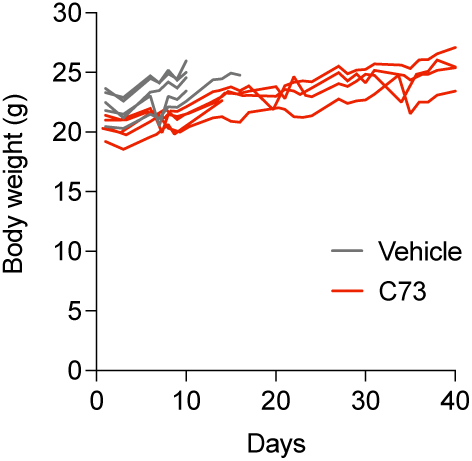
Mice body weight over *in vivo* experiment.

## Notes

### Competing Interest Statement

The authors have declared no competing interest.

## REFERENCES

1. Tang, H. et al. Fusion genes in cancers: Biogenesis, functions, and therapeutic implications. Genes & Diseases, 101536 (2025).

2. Ren, R. Mechanisms of BCR–ABL in the pathogenesis of chronic myelogenous leukaemia. Nature Reviews Cancer 5, 172–183 (2005).

3. Delattre, O. et al. Gene fusion with an ETS DNA-binding domain caused by chromosome translocation in human tumours. Nature 359, 162–165 (1992).

4. Liu, S. V., Nagasaka, M., Atz, J., Solca, F. & Müllauer, L. Oncogenic gene fusions in cancer: from biology to therapy. Signal Transduction and Targeted Therapy 10, 111 (2025).

5. Bushweller, J. H. Targeting transcription factors in cancer – from undruggable to reality. Nat Rev Cancer 19, 611–624 (2019).

6. Jawad, M. U. et al. Ewing sarcoma demonstrates racial disparities in incidence-related and sex-related differences in outcome. Cancer 115, 3526–3536 (2009).

7. Gaspar, N. et al. Ewing Sarcoma: Current Management and Future Approaches Through Collaboration. J Clin Oncol 33, 3036–3046 (2015).

8. Stahl, M. et al. Risk of recurrence and survival after relapse in patients with Ewing sarcoma. Pediatr Blood Cancer 57, 549–553 (2011).

9. Grünewald, T. G. P. et al. Ewing sarcoma. Nat Rev Dis Primers 4, 5 (2018).

10. Ohno, T., Rao, V. N. & Reddy, E. S. EWS/Fli-1 chimeric protein is a transcriptional activator. Cancer Res 53, 5859–5863 (1993).

11. Riggi, N. et al. EWS-FLI1 utilizes divergent chromatin remodeling mechanisms to directly activate or repress enhancer elements in Ewing sarcoma. Cancer Cell 26, 668–681 (2014).

12. Flores, G. & Grohar, P. J. One oncogene, several vulnerabilities: EWS/FLI targeted therapies for Ewing sarcoma. Journal of Bone Oncology 31, 100404 (2021).

13. Yasir, M., Park, J. & Chun, W. EWS/FLI1 Characterization, Activation, Repression, Target Genes and Therapeutic Opportunities in Ewing Sarcoma. Int J Mol Sci 24 (2023).

14. Potikyan, G. et al. Genetically defined EWS/FLI1 model system suggests mesenchymal origin of Ewing’s family tumors. Lab Invest 88, 1291–1302 (2008).

15. Lessnick, S. L., Dacwag, C. S. & Golub, T. R. The Ewing’s sarcoma oncoprotein EWS/FLI induces a p53-dependent growth arrest in primary human fibroblasts. Cancer Cell 1, 393–401 (2002).

16. Deneen, B. & Denny, C. T. Loss of p16 pathways stabilizes EWS/FLI1 expression and complements EWS/FLI1 mediated transformation. Oncogene 20, 6731–6741 (2001).

17. Sohn, E. J. et al. EWS/FLI1 oncogene activates caspase 3 transcription and triggers apoptosis in vivo. Cancer Res 70, 1154–1163 (2010).

18. Minas, T. Z. et al. Combined experience of six independent laboratories attempting to create an Ewing sarcoma mouse model. Oncotarget 8, 34141–34163 (2017).

19. Seong, B. K. A. et al. TRIM8 modulates the EWS/FLI oncoprotein to promote survival in Ewing sarcoma. Cancer Cell 39, 1262–1278.e1267 (2021).

20. Nieto-Soler, M. et al. Efficacy of ATR inhibitors as single agents in Ewing sarcoma. Oncotarget 7, 58759–58767 (2016).

21. Su, X. A. et al. RAD21 is a driver of chromosome 8 gain in Ewing sarcoma to mitigate replication stress. Genes Dev 35, 556–572 (2021).

22. Gorthi, A. et al. EWS-FLI1 increases transcription to cause R-loops and block BRCA1 repair in Ewing sarcoma. Nature 555, 387–391 (2018).

23. Zeman, M. K. & Cimprich, K. A. Causes and consequences of replication stress. Nat Cell Biol 16, 2–9 (2014).

24. Musa, J. et al. Cooperation of cancer drivers with regulatory germline variants shapes clinical outcomes. Nat Commun 10, 4128 (2019).

25. Zhou, Y. et al. Pan-Cancer Analysis of Oncogenic Role of RAD54L and Experimental Validation in Hepatocellular Carcinoma. J Inflamm Res 16, 3997–4017 (2023).

26. Keijzers, G. et al. Human Exonuclease 1 (EXO1) Regulatory Functions in DNA Replication with Putative Roles in Cancer. Int J Mol Sci 20 (2018).

27. Mackintosh, C. et al. 1q gain and CDT2 overexpression underlie an aggressive and highly proliferative form of Ewing sarcoma. Oncogene 31, 1287–1298 (2012).

28. Díaz-Martín, J. et al. Prospective evaluation of copy number alterations validates chromosome 1q gain as an independent marker of poor prognosis in localized Ewing sarcoma. Experimental and Molecular Pathology 144, 105008 (2025).

29. Tirode, F. et al. Genomic landscape of Ewing sarcoma defines an aggressive subtype with co-association of STAG2 and TP53 mutations. Cancer Discov 4, 1342–1353 (2014).

30. García-Domínguez, D. J. et al. An inducible ectopic expression system of EWSR1-FLI1 as a tool for understanding Ewing sarcoma oncogenesis. PLoS One 15, e0234243 (2020).

31. Paiano, J. et al. Role of 53BP1 in end protection and DNA synthesis at DNA breaks. Genes Dev 35, 1356–1367 (2021).

32. Rogakou, E. P., Boon, C., Redon, C. & Bonner, W. M. Megabase chromatin domains involved in DNA double-strand breaks in vivo. J Cell Biol 146, 905–916 (1999).

33. Gnügge, R. & Symington, L. S. DNA end resection during homologous recombination. Curr Opin Genet Dev 71, 99–105 (2021).

34. van de Kooij, B. et al. EXO1 protects BRCA1-deficient cells against toxic DNA lesions. Mol Cell 84, 659–674.e657 (2024).

35. Xie, Y. et al. RBX1 prompts degradation of EXO1 to limit the homologous recombination pathway of DNA double-strand break repair in G1 phase. Cell Death & Differentiation 27 (2019).

36. Fernandez-Vidal, A., Vignard, J. & Mirey, G. Around and beyond 53BP1 Nuclear Bodies. Int J Mol Sci 18 (2017).

37. Krupina, K., Goginashvili, A. & Cleveland, D. W. Causes and consequences of micronuclei. Curr Opin Cell Biol 70, 91–99 (2021).

38. Carrillo, J. et al. Cholecystokinin down-regulation by RNA interference impairs Ewing tumor growth. Clin Cancer Res 13, 2429–2440 (2007).

39. Fragkos, M. & Naim, V. Rescue from replication stress during mitosis. Cell Cycle 16, 613–633 (2017).

40. Bertolin, A. P., Hoffmann, J. S. & Gottifredi, V. Under-Replicated DNA: The Byproduct of Large Genomes? Cancers (Basel) 12 (2020).

41. Baumann, C., Körner, R., Hofmann, K. & Nigg, E. A. PICH, a centromere-associated SNF2 family ATPase, is regulated by Plk1 and required for the spindle checkpoint. Cell 128, 101–114 (2007).

42. Bhowmick, R., Minocherhomji, S. & Hickson, I. D. RAD52 Facilitates Mitotic DNA Synthesis Following Replication Stress. Molecular Cell 64, 1117–1126 (2016).

43. Chan, Y. W., Fugger, K. & West, S. C. Unresolved recombination intermediates lead to ultra-fine anaphase bridges, chromosome breaks and aberrations. Nature Cell Biology 20, 92–103 (2018).

44. Yoshida, A. et al. NKX2.2 is a Useful Immunohistochemical Marker for Ewing Sarcoma. The American Journal of Surgical Pathology 36, 993–999 (2012).

45. Zöllner, S. K. et al. Ewing Sarcoma-Diagnosis, Treatment, Clinical Challenges and Future Perspectives. J Clin Med 10 (2021).

46. García-Rodríguez, N., Domínguez-García, I., Domínguez-Pérez, M. del C. & Huertas, P. EXO1 and DNA2-mediated ssDNA gap expansion is essential for ATR activation and to maintain viability in BRCA1-deficient cells. Nucleic Acids Research 52, 6376–6391 (2024).

47. Yang, G. et al. EXO1 Plays a Carcinogenic Role in Hepatocellular Carcinoma and is related to the regulation of FOXP3. Journal of Cancer 11, 4917–4932 (2020).

48. Wang, Z. et al. EXO1/P53/SREBP1 axis-regulated lipid metabolism promotes prostate cancer progression. Journal of Translational Medicine 22, 104 (2024).

49. Bolderson, E. et al. Phosphorylation of EXO1 modulates homologous recombination repair of DNA double-strand breaks. Nucleic acids research 38, 1821–1831 (2009).

50. Gao, Y. et al. A CRISPR-Cas9 screen identifies EXO1 as a formaldehyde resistance gene. Nature Communications 14, 381 (2023).

51. Yang, H. et al. Cyclin F–EXO1 axis controls cell cycle–dependent execution of double-strand break repair. Science Advances 10, eado0636

52. Stroik, S. et al. EXO1 resection at G-quadruplex structures facilitates resolution and replication. Nucleic acids research 48 (2020).

53. Mocanu, C. & Chan, K. L. Mind the replication gap. R Soc Open Sci 8, 201932 (2021).

54. Audrey, A., et al. RAD52-dependent mitotic DNA synthesis is required for genome stability in Cyclin E1-overexpressing cells. Cell Reports 43 (2024).

55. Mocanu, C. et al. DNA replication is highly resilient and persistent under the challenge of mild replication stress. Cell Rep 39, 110701 (2022).

56. Tomimatsu, N. et al. DNA-damage-induced degradation of EXO1 exonuclease limits DNA end resection to ensure accurate DNA repair. J Biol Chem 292, 10779–10790 (2017).

57. Lydeard, J. R., Lipkin-Moore, Z., Jain, S., Eapen, V. V. & Haber, J. E. Sgs1 and Exo1 Redundantly Inhibit Break-Induced Replication and De Novo Telomere Addition at Broken Chromosome Ends. PLOS Genetics 6, e1000973 (2010).

58. Marrero, V. A. & Symington, L. S. Extensive DNA End Processing by Exo1 and Sgs1 Inhibits Break-Induced Replication. PLOS Genetics 6, e1001007 (2010).

59. Barwacz, S. A. et al. DNA double-strand break end resection factors and WRN facilitate mitotic DNA synthesis in human cells. Nat Commun 16, 7901 (2025).

60. Wang, Y. et al. Discovery and Characterization of Small Molecule Inhibitors Targeting Exonuclease 1 for Homologous Recombination-Deficient Cancer Therapy. ACS Chem Biol 20, 1258–1272 (2025).

61. Zhou, C. S. et al. Exonuclease 1 (EXO1) is a Potential Prognostic Biomarker and Correlates with Immune Infiltrates in Lung Adenocarcinoma. Onco Targets Ther 14, 1033–1048 (2021).

62. Dai, Y. et al. EXO1 overexpression is associated with poor prognosis of hepatocellular carcinoma patients. Cell Cycle 17, 2386–2397 (2018).

63. Luo, F., Wang, Y. Z., Lin, D., Li, J. & Yang, K. Exonuclease 1 expression is associated with clinical progression, metastasis, and survival prognosis of prostate cancer. J Cell Biochem 120, 11383–11389 (2019).

64. Wiederschain, D. et al. Single-vector inducible lentiviral RNAi system for oncology target validation. Cell Cycle 8, 498–504 (2009).

65. Olmedo-Pelayo, J. et al. EWS::FLI1-DHX9 interaction promotes Ewing sarcoma sensitivity to DNA topoisomerase 1 poisons by altering R-loop metabolism. Oncogene, 44, 3537–3552 (2025).

66. Puerto-Camacho, P. et al. Preclinical Efficacy of Endoglin-Targeting Antibody-Drug Conjugates for the Treatment of Ewing Sarcoma. Clin Cancer Res 25, 2228–2240 (2019).

67. Orth, M. F. et al. Systematic multi-omics cell line profiling uncovers principles of Ewing sarcoma fusion oncogene-mediated gene regulation. Cell Rep 41, 111761 (2022).

68. Reich, M. et al. in Nat Genet Vol. 38 500–501 (2006).

